# Ferroportin 3 is a dual-targeted mitochondrial/chloroplast iron exporter necessary for iron homeostasis in Arabidopsis

**DOI:** 10.1101/2020.07.15.203646

**Authors:** Leah J. Kim, Kaitlyn M. Tsuyuki, Fengling Hu, Emily Y. Park, Jingwen Zhang, Jennifer Gallegos Iraheta, Ju-Chen Chia, Rong Huang, Avery E. Tucker, Madeline Clyne, Claire Castellano, Angie Kim, Daniel D. Chung, Christopher T. DaVeiga, Elizabeth M. Parsons, Olena K. Vatamaniuk, Jeeyon Jeong

## Abstract

Mitochondria and chloroplasts are organelles with high iron demand that are particularly susceptible to iron-induced oxidative stress. Despite the necessity of strict iron regulation in these organelles, much remains unknown about mitochondrial and chloroplast iron transport in plants. Here, we propose that Arabidopsis Ferroportin 3 (FPN3) is an iron exporter dual-targeted to mitochondria and chloroplasts. *FPN3* is expressed in shoots regardless of iron conditions, but its transcripts accumulate under iron deficiency in roots. *fpn3* mutants cannot grow as well as wild type under iron-deficient conditions and shoot iron levels are reduced in *fpn3* mutants compared to wild type. ICP-MS measurements show that iron levels in the mitochondria and chloroplasts are increased relative to wild type, consistent with the proposed role of FPN3 as a mitochondrial/plastid iron exporter. In iron deficient *fpn3* mutants, abnormal mitochondrial ultrastructure was observed, whereas chloroplast ultrastructure was not affected, implying that FPN3 plays a critical role in the mitochondria. Overall, our study suggests that FPN3 is essential for optimal iron homeostasis.

**Significance statement:** Iron homeostasis must be tightly controlled in the mitochondria and chloroplasts, but iron trafficking in these organelles is not fully understood. Our work suggests that FPN3 is an iron exporter required for maintaining proper iron levels in mitochondria and chloroplasts. Furthermore, FPN3 is necessary for the optimal growth and normal mitochondrial ultrastructure under iron deficiency. This study reveals the physiological role of FPN3 and advances our understanding of iron regulation in mitochondria and chloroplasts.

## INTRODUCTION

Iron serves as a critical redox cofactor in vital cellular processes. Nevertheless, excess or improperly regulated iron can cause deleterious effects by generating hydroxyl radicals via the Fenton reaction (Halliwell and Gutteridge, 1992). Therefore, iron homeostasis must be tightly maintained in all organisms, including plants. As photosynthetic organisms, plants use iron as an essential cofactor in both respiration and photosynthesis. At the same time, plant cells must regulate iron to ensure adequate supply while avoiding oxidative stress (Shcolnick and Keren, 2006). Despite being an abundant element in the soil, iron is one of the most limiting nutrients for plant growth – it has extremely low bioavailability under aerobic conditions at neutral or alkaline pH (Marschner, 2012; Colombo *et al.*, 2014). Iron homeostasis in plants is of particular interest, because understanding its mechanisms will provide insights to improving agriculture and human health (Vasconcelos *et al.*, 2017).

Dicots acquire iron by a reduction-based mechanism that is induced under iron deficiency (Jeong *et al.*, 2017; Connorton *et al.*, 2017; Kobayashi *et al.*, 2018; Brumbarova *et al.*, 2015). Ferric chelates in the rhizosphere are solubilized by the release of protons (Santi and Schmidt, 2009) and reduced to ferrous iron by FERRIC REDUCTASE OXIDASE 2 (FRO2) (Robinson *et al.*, 1999). Coumarins released from iron-deficient roots are considered to aid this process as well (Clemens and Weber, 2016; Schmidt *et al.*, 2014; Fourcroy *et al.*, 2014). Ferrous iron is then transported into the root via IRON-REGULATED TRANSPORTER 1 (IRT1) (Connolly *et al.*, 2002; Vert *et al.*, 2002; Varotto *et al.*, 2002). Once iron reaches the root vasculature, it is loaded into the xylem by FERROPORTIN 1 (FPN1) (Morrissey *et al.*, 2009) and chelated with citrate, which is transported into the xylem by the FERRIC REDUCTASE DEFECTIVE 3 (FRD3) transporter (Durrett *et al.*, 2007). The iron-citrate complex is then translocated to shoots. For lateral translocation in shoots, iron-nicotianamine (NA) complexes are formed and translocated from leaves to seeds via the phloem by YELLOW STRIPE-LIKE (YSL) family members (DiDonato *et al.*, 2004; Schaaf *et al.*, 2005; Waters *et al.*, 2006). Two YSL members, YSL1 and YSL3, were shown to regulate long-distance iron-deficiency signals from shoots (Kumar *et al.*, 2017) and are responsible for loading iron into the seeds (Le Jean *et al.*, 2005; Waters *et al.*, 2006). Oligopeptide transporter 3 (OPT3) also plays a role in transmitting shoot-to-root iron signals and regulates the redistribution of iron from source to sink tissues, such as from old leaves to seeds or to developing tissues (Mendoza-Cózatl *et al.*, 2014; Zhai *et al.*, 2014; Stacey *et al.*, 2008).

In addition to iron acquisition and its distribution between tissues, iron trafficking across subcellular compartments is crucial for proper iron homeostasis. In particular, chloroplasts and mitochondria need a substantial amount of iron; many components of the photosynthetic and respiratory electron transport chains use iron as a cofactor. Fe-S cluster assembly occurs in these organelles and heme biosynthesis occurs in plastids (Masuda *et al.*, 2003; Tanaka *et al.*, 2011). Proteins involved in the last stage of heme biosynthesis are also present in mitochondria (Balk and Schaedler, 2014). Chloroplasts are the most iron-rich organelle in plant cells and accounts for 60-80% of iron found in a leaf cell (Terry and Low, 1982; Shikanai *et al.*, 2003). Meanwhile, in the mitochondria, iron is the major micronutrient present with a molar ratio of 26:8:6:1 for Fe:Zn:Cu:Mn (Tan *et al.*, 2010). Mitochondria and chloroplasts are also highly susceptible to oxidative stress due to reactive oxygen species (ROS) generated by the electron transport chain. It has been shown that the iron sequestering protein ferritin is present in mitochondria and plastids, and helps to prevent iron-induced oxidative stress (Briat *et al.*, 2010; Zancani *et al.*, 2004; Tarantino, Casagrande, *et al.*, 2010). In the mitochondria, frataxin, which is involved in Fe-S cluster and heme biogenesis, also plays a role in protection against iron-induced oxidative stress (Gomez-Casati *et al.*, 2018).

Iron transport in mitochondria and chloroplasts is not as well understood as the mechanisms of iron acquisition in the roots, but molecular and physiological studies have been gradually contributing to understanding iron regulation in chloroplasts and mitochondria. In chloroplasts, physiological studies suggest that ferric chelates move across the outer membrane (Bughio *et al.*, 1997; Solti *et al.*, 2012; Müller *et al.*, 2019), but ferrous iron is imported into chloroplasts across the inner membrane (Bughio *et al.*, 1997; Shingles *et al.*, 2002). The ferric chelate reductase, FRO7, reduces iron for chloroplast iron acquisition (Jeong *et al.*, 2008), and PERMEASE IN CHLOROPLAST 1 (PIC1) mediates iron transport into chloroplasts via interaction with NiCo (Duy *et al.*, 2007; Duy *et al.*, 2011). A recent study in *Brassica napus* proposes that NiCo may also be involved in iron sensing or iron release from chloroplasts (Pham *et al.*, 2020). The Arabidopsis MitoFerrinLike1 (Mfl1) has also been reported to import iron into chloroplasts (Tarantino *et al.*, 2011). In addition, ATP-binding cassette transporters, ABCI10, NAP14/ABCI11, and ABCI12, may also contribute to iron uptake into chloroplasts (Shimoni-Shor *et al.*, 2010; Voith von Voithenberg *et al.*, 2019). YSL4 and YSL6 have been reported to efflux iron-NA complexes from the chloroplasts (Divol *et al.*, 2013). However, both transporters were also identified in the tonoplast proteome (Jaquinod *et al.*, 2007) and were targeted to the vacuolar and intracellular membranes (Conte *et al.*, 2013). ZmFRD4 was reported as a potential thylakoid iron importer in maize, although direct evidence for iron transport remains to be found (Zhang *et al.*, 2017).

In the mitochondria, a mitoferrin ortholog from rice, MITOCHONDRIAL IRON TRANSPORTER 1 (MIT1), was identified as a mitochondrial iron importer (Bashir *et al.*, 2011). Two Arabidopsis mitoferrin orthologs, MIT1 and MIT2, have recently been reported as mitochondrial iron importers that mediate cellular iron homeostasis and are essential for embryogenesis (Jain *et al.*, 2019). A reduction-based mechanism might be involved in moving iron in and out of mitochondria, because ferric chelate reductases, FRO3 and FRO8, are localized to the mitochondria in Arabidopsis (Jeong and Connolly, 2009; Heazlewood *et al.*, 2004). The two mitochondrial FROs are likely to play non-overlapping roles, as *FRO3* is induced by iron deficiency, whereas *FRO8* is not iron-regulated (Jeong and Connolly, 2009; Mukherjee *et al.*, 2006).

In this study, we investigated an Arabidopsis FPN family member, FPN3/IREG3. FPNs, also known as solute carrier (SCL) group 40A1 and IRON REGULATED (IREG) transporters, efflux iron from the cytoplasm (Drakesmith *et al.*, 2015). In animals, FPNs are found in cells that play critical roles in supplying iron, such as intestinal cells. While animal species have only one FPN, multiple FPN paralogs are present within most plant species. In Arabidopsis, three FPN/IREG family members were identified based on phylogenetic analysis (Schaaf et al., 2006). FPN1/IREG1 (At2g38460) and FPN2/IREG2 (At5g03570) share 77 % identity, whereas FPN3/IREG3 (At5g26820) shares about 20% identity with FPN1 and FPN2 (Figure 1). FPN1/IREG1 localizes to the plasma membrane of cells in the stele, and FPN2/IREG2 is localized on the vacuolar membrane to buffer cellular metal influx (Morrissey *et al.*, 2009). Previously, FPN3/IREG3 was reported as Multiple Antibiotic Resistance 1 (MAR1), a chloroplast protein that allows antibiotics to opportunistically enter chloroplasts (Conte *et al.*, 2009). While it was speculated that MAR1 may be involved in iron homeostasis, its potential role in iron regulation and its physiological function were not previously studied. Here, we report that FPN3 is dual-targeted to mitochondria and chloroplasts, and provide results suggesting that FPN3 exports iron from these organelles. Furthermore, our study provides key evidence indicating that FPN3 plays a critical role in iron homeostasis and its function is important in the mitochondria as evidenced by the drastic morphological changes in *fpn3* mitochondria under iron deficient conditions.

**Figure 1.**
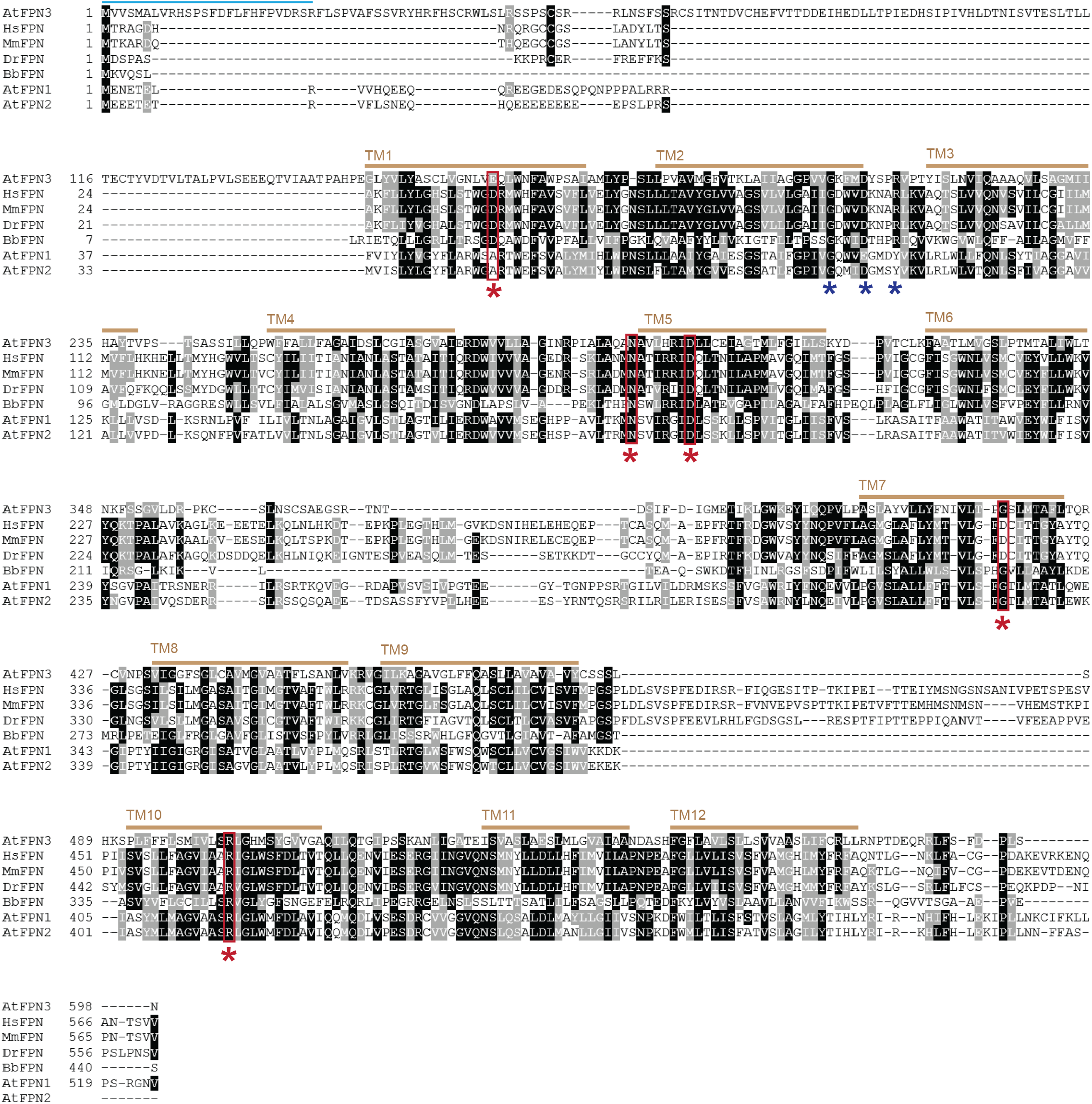
Sequence alignment of FPNs. Multiple sequence alignment was performed using T-Coffee (Notredame *et al.*, 2000) with FPN amino acid sequences of *Arabidopsis thaliana* (AtFPN1, AtFPN2, AtFPN3), *Homo sapiens* (HsFPN), *Mus musculus* (MmFPN), *Danio reio* (DrFPN), and *Bdellovibrio bacteriovorus* (BbFPN). Identical residues are shaded in black, whereas similar residues are shaded in grey. Red stars indicate residues involved in iron binding and transport in human and bacterial FPNs (Bonaccorsi di Patti *et al.*, 2015; Taniguchi *et al.*, 2015). The blue stars indicate motif A, GX3DX3R, which is conserved in MFS members. The predicted transit peptide, denoted as a blue bar, suggests a chloroplast or mitochondrial localization (Schwacke *et al.*, 2003). The yellow brown bars indicate predicted transmembrane domains.

## RESULTS

### Iron binding and transport residues of human FPN are conserved in Arabidopsis FPN3

To investigate the potential role of FPN3 in iron homeostasis, we first examined if conserved iron binding and transport residues of other FPN members are present in FPN3. Structural modeling and *in vitro* assays with mutated variants of human FPN identified amino acids that are critical for iron binding and transport (Bonaccorsi di Patti *et al.*, 2014; Taniguchi *et al.*, 2015). These residues were highly conserved across FPN orthologs in human, mouse, zebrafish, frog, bacteria, and Arabidopsis FPN paralogs (Taniguchi *et al.*, 2015). For example, two Asp residues of the human FPN, Asp39 and Asp181, that are essential for iron transport (Bonaccorsi di Patti *et al.*, 2014; Taniguchi *et al.*, 2015) were highly conserved in FPN3 as indicated by presence of the Glu, which is synonymous to Asp, and Asp residues in their corresponding positions (Figure 1). Two other iron binding/transport residues of HsFPN, Asp325 and Asn174, were conserved in FPN3 as well (Figure 1). Arg466 of HsFPN, another amino acid that affected iron efflux (Bonaccorsi di Patti *et al.*, 2014; Taniguchi *et al.*, 2015), was not preserved in FPN3 (Figure 1). However, this Arg is not part of the metal binding site (Bonaccorsi di Patti *et al.*, 2014; Taniguchi *et al.*, 2015) and the Gly found in this position was conserved among the bacterial and Arabidopsis FPNs. Overall, the conserved iron binding/transport residues suggest that FPN3 is highly likely to transport ferrous iron, which is a major substrate of FPN orthologs(Drakesmith *et al.*, 2015; Taniguchi *et al.*, 2015).

### Heterologously expressed *FPN3* exports iron from mitochondria in yeast

Multiple chloroplast proteins have been targeted to the mitochondria when expressed in fungi (Versaw and Harrison, 2002; Jeong *et al.*, 2008; Hurt *et al.*, 1986; Pfaller *et al.*, 1989; Brink *et al.*, 1994). Based on the prior report on MAR1 that indicated chloroplast localization (Conte *et al.*, 2009) and the predicted transit peptides that suggested targeting to chloroplasts or mitochondria (Schwacke *et al.*, 2003), we tested if FPN3 would be able to complement mitochondrial iron transporter mutants of *Saccharomyces cerevisiae*. After verifying that FPN3 is localized to mitochondria using Western blots with mitochondrial fractions of yeast cells expressing FPN3-FLAG (Supplementary Figure 1), we expressed *FPN3* in Δ*mmt1/2*, which lack mitochondrial iron exporters. Previous studies showed that Δ*mmt1/2* did not have a strong phenotype, but the deletion of *MMT1/2* in a Δ*ccc1* background, which lacks a vacuolar iron importer, resulted in a slightly decreased sensitivity to high iron compared to Δ*ccc1* (Li *et al.*, 2014). Additionally, Δ*ccc1*Δ*mmt1/2* cells expressing *MMT1/2* are highly sensitive to high iron due to increased cytosolic iron (Li *et al.*, 2014). In the present study, we expressed *FPN3* in Δ*ccc1*Δ*mmt1/2* under high iron conditions to test if phenotypes similar to Δ*ccc1*Δ*mmt1/2* cells expressing *MMT1/2* would arise. Consistent with observations by Li et al. (2014), the expression of *MMT1/2* resulted in reduced growth in high iron (Figure 2A, 2B). The growth of Δ*ccc1*Δ*mmt1/2* expressing *FPN3* was decreased in high iron and approximated that of Δ*ccc1*Δ*mmt1/2* expressing *MMT1/2* (Figure 2A, 2B). Thus, our high iron growth assay results from both plates and liquid cultures suggest that FPN3 exports iron from the mitochondria (Figure 2A, 2B).

**Figure 2.**
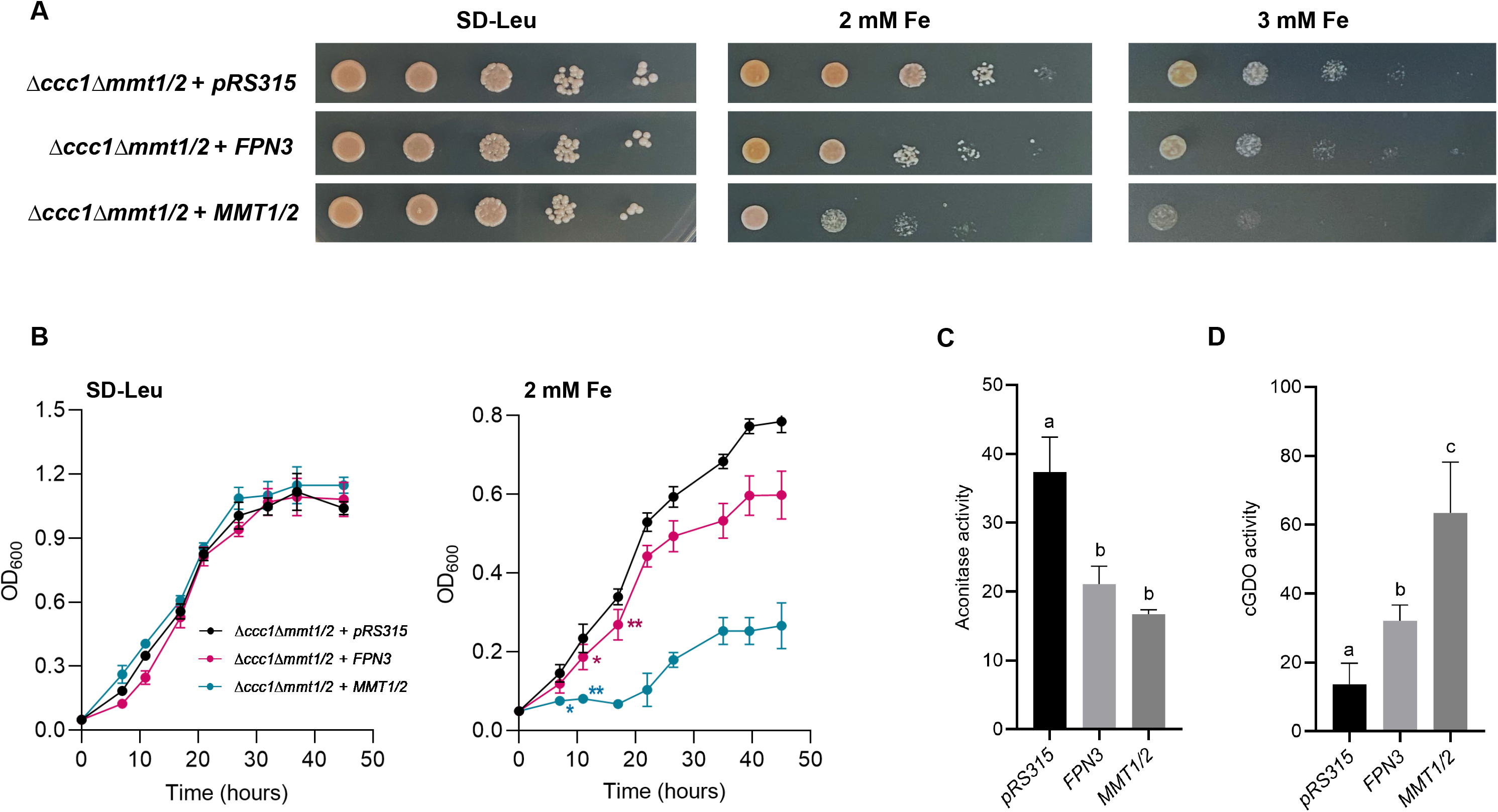
Heterologous expression of *FPN3* in yeast. (A) Spot assays in high iron media (SD-Leu with 2 mM and 3 mM Fe) or control media (SD-Leu) with *Δccct1Δmmt1/2* cells expressing *FPN3* or *MMT1/2*, or transformed with an empty vector, *pRS315*. (B) Growth assays with liquid culture with *Δccct1Δmmt1/2* cells expressing *FPN3* or *MMT1/2*, or transformed with an empty vector, *pRS315*. Significant differences compared to the negative control were determined by two-way ANOVA followed by Tukey’s multiple comparison test (*: p<0.05; **: p<0.001; n=6; Error bars = SD). From the third time point and beyond, cells expressing *FPN3* or *MMT1/2* were significantly different from the negative control (p<0.001) but labels were omitted for simplicity of the graph. (C) Aconitase activity (nmol/mg protein/min) of wild type (DY150) cells expressing *FPN3* or *MMT1/2* or transformed with an empty vector, *pRS426* (n=6; Error bars = SD). (D) cGDO activity (mmol/mg protein/min) of wild type (DY150) cells expressing *FPN3*, *MMT1/2,* or the empty vector, *pRS315.* Significant differences were determined by one-way ANOVA followed by Tukey’s test (*: p<0.01; n=6; Error bars = SD).

We also expressed *FPN3* in the *Δmrs3Δmrs4* yeast mutant, which lacks the mitochondrial iron importers, Mrs3 and Mrs4, to test if FPN3 is likely to import iron into mitochondria. The *Δmrs3Δmrs4* mutant does not have a strong iron phenotype, but exhibits increased sensitivity to oxidative stress compared to wild type cells (Mühlenhoff *et al.*, 2003; Foury and Roganti, 2002). Expression of *FPN3* in *Δmrs3Δmrs4* did not rescue the oxidative stress sensitivity of *Δmrs3Δmrs4* (Supplementary Figure 2), which would be expected if *FPN3* were a mitochondrial iron importer. By contrast, *FPN3* expression exacerbated the phenotype, consistent with that of cells expressing *MMT1/2* (Supplementary Figure 2), indicating that FPN3 is not likely to be importing iron into the mitochondria.

Based on the results from Δ*ccc1*Δ*mmt1/2* and *Δmrs3Δmrs4* cells expressing *FPN3* or *MMT1/2*, we hypothesized that FPN3 is a mitochondrial iron exporter. If FPN3 is exporting iron from the mitochondria, mitochondrial iron levels should be lower in cells overexpressing *FPN3*. To test this hypothesis, we measured the activity of aconitase in mitochondria to indirectly assess mitochondrial iron levels. Aconitase is a Fe-S enzyme involved in the tricarboxylic acid cycle, and its activity correlates with the availability of mitochondrial iron. For example, decreased aconitase activity has been detected in the yeast *Δmrs3/4* cells and rice plants defective in mitochondrial iron import (Foury and Roganti, 2002; Bashir *et al.*, 2011). Although aconitase is present in the mitochondria and the cytosol, mitochondrial aconitase accounts for most of the cellular aconitase activity in yeast, and yeast cytosolic aconitase is not involved in regulating iron metabolism unlike in mammalian cells (Regev-Rudzki *et al.*, 2005). Aconitase activity of cells expressing *FPN3* was similar to cells expressing *MMT1/2* and exhibited about 30% lower activity than that of the negative control with an empty vector (Figure 2C), suggesting that FPN3 is exporting iron from the mitochondria.

As an alternative approach to testing mitochondrial iron export by FPN3, we examined if cytosolic iron levels increased in cells expressing *FPN3*. We co-transformed *FPN3* or control plasmids along with a plasmid that expresses the bacterial gentisate 1,2-dioxygenase (GDO) in the yeast cytoplasm (Li *et al.*, 2012) and measured GDO activity. GDO uses iron as a cofactor and the activity of cytosolic GDO (c-GDO), which has been confirmed to localize to the cytoplasm in yeast, correlates with the amount of cytosolic iron. Thus, c-GDO can serve as an indicator of cytosolic iron levels in yeast (Li *et al.*, 2012; Li *et al.*, 2014). We detected significantly higher c-GDO activity in cells expressing *FPN3* or *MMT1/2*, whereas only background level activity was observed in cells with the empty vector control (Figure 2D). This result is consistent with our findings from yeast growth tests and aconitase activity assays (Figure 2A-C). Overall, our yeast results consistently supported the hypothesis that FPN3 is a mitochondrial iron exporter and is functionally analogous to Mmt1/2.

### *FPN3* is expressed in the shoots, roots, flowers, and siliques, and is iron-regulated in roots

To detect the expression of *FPN3* at the tissue level, we fused the promoter region of *FPN3* to the β-glucuronidase (GUS) reporter gene (*FPN3p-GUS*), transformed the construct into wild type Arabidopsis, and conducted GUS histochemical staining. *FPN3p-GUS* was expressed in shoots and roots of seedlings from an early stage, and in floral organs and siliques (Figure 3A-F), which was in agreement with *FPN3* expression data reported in multiple transcriptomics studies (Schmid *et al.*, 2005; Zimmermann *et al.*, 2004; Dinneny *et al.*, 2008; Winter *et al.*, 2007). To determine whether *FPN3* expression is regulated by iron, we performed GUS staining with *FPN3p-GUS* plants grown under iron-deficient or iron-sufficient conditions. We detected stronger staining in roots of seedlings from iron-deficient medium than in roots from iron-sufficient medium, whereas *FPN3p-GUS* staining in the shoots was prominent under both conditions (Figure 3I-K). *FPN3p-GUS* expression in 1-day old seedlings exhibited similar patterns, with more intense staining in the roots of plants grown under iron-deficient conditions compared to plants from iron-sufficient conditions (Figure 3G, H). Staining in the shoots was similar under both iron conditions. This observation corroborates our RT-qPCR results in which *FPN3* transcript level was approximately 4.5-fold higher in iron-deficient roots (Figure 3L), and is consistent with transcriptomics studies that reported higher steady state levels of *FPN3* transcripts in iron-deficient roots (Mai *et al.*, 2016; Dinneny *et al.*, 2008; Yang *et al.*, 2010; Park *et al.*, 2019; Khan *et al.*, 2018; Buckhout *et al.*, 2009). Meanwhile, the constitutive expression of *FPN3* was detected in the shoots regardless of the iron status of the plant (Figure 3I, L).

**Figure 3.**
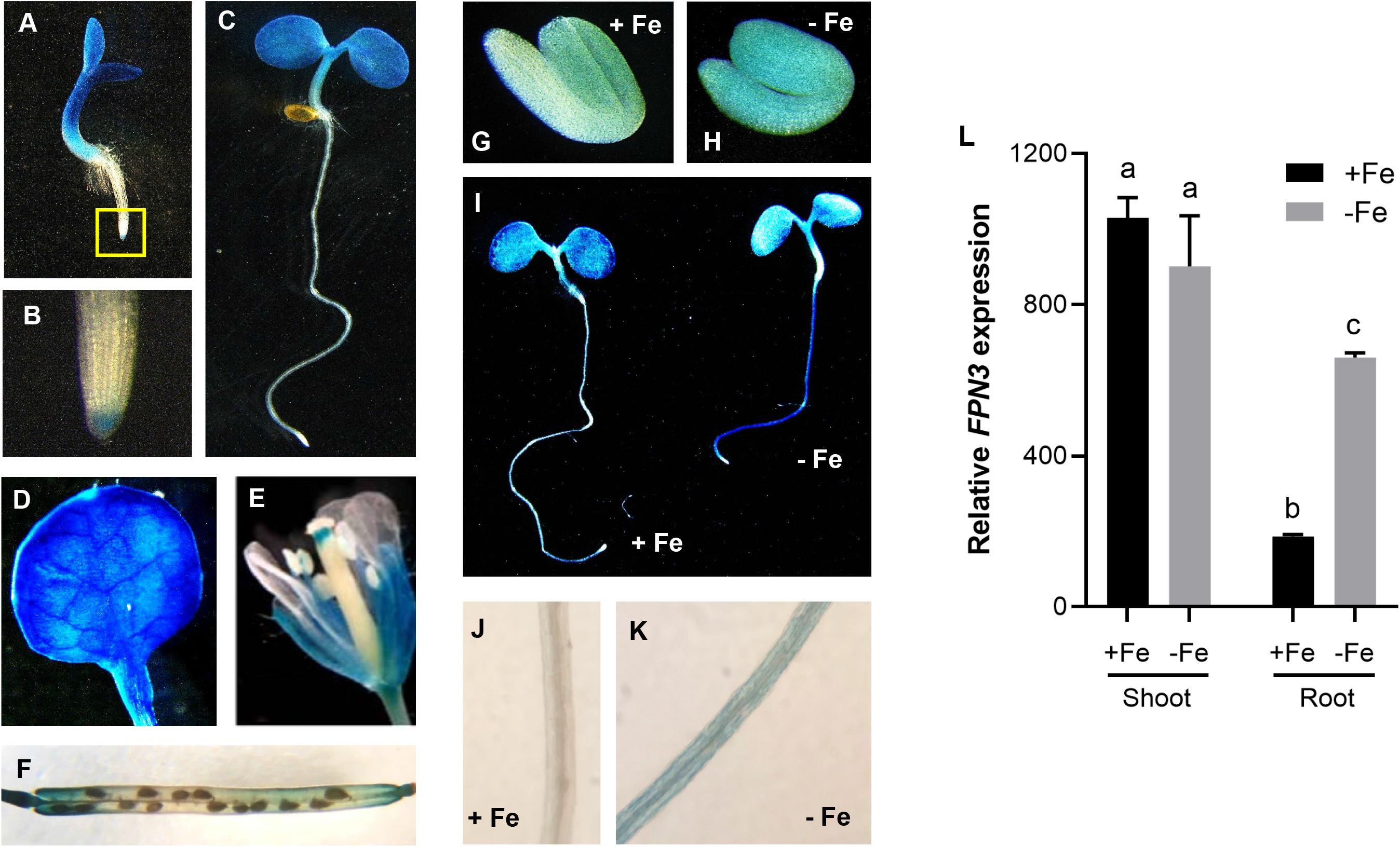
*FPN3* expression detected by *FPN3p-GUS* and RT-qPCR. (A-C) Histochemical staining of *FPN3p-GUS* seedlings germinated under iron sufficient conditions on day 3 (A) and day 5 (C), and a close-up image of the root tip of a day 3 seedling (B; boxed region in A). (D) *FPN3p-GUS* expression in the first true leaf of a 3-week old plant. (E, F) *FPN3p-GUS* staining in floral organs (E) and the silique (F). (G, H) *FPN3p-GUS* staining in 1-day-old imbibing seedlings germinated on iron deficient (G) or sufficient (H) media. (I) *FPN3p-GUS* staining in 3-day-old seedlings germinated on iron deficient (−Fe) or sufficient (+Fe) media. (J, K) Close-up view of iron sufficient root (J) and iron deficient root (K). Representative images of seedlings or organs from at least 6 individuals are shown. (L) Relative expression of *FPN3* in shoots and roots of wild type plants treated under iron deficient or sufficient conditions. *FPN3* transcript level was quantified by RT-qPCR and normalized with *ACT2.* (ANOVA; Tukey’s test; *: p<0.05; n=3; error bars = SD).

### *FPN3* is iron-regulated primarily by local signals in roots

Studies with multiple plant species have revealed that two types of iron-deficiency signals regulate the iron-deficiency response in the roots: the local signal determined by the iron level in the rhizosphere, and the systemic, long distance signal based on the iron status of the shoots (Vert *et al.*, 2003; Grusak and Pezeshgi, 1996; García *et al.*, 2013; Bienfait *et al.*, 1987). The differential level of *FPN3* transcripts in roots and shoots prompted us to test if the iron-dependent *FPN3* accumulation in roots is regulated by a local or long-distance signal. Thus, we generated two lateral roots, or split roots, and each root was subjected to different iron growth conditions. The iron growth conditions for our split-root experiments were optimized based on the method by Kumar et al. (2017). Plants were grown under iron-deficient or iron-sufficient conditions for 3 days prior to the split media experiment and RNA was extracted from each side of the split root after 0, 6, or 12 hours after transfer (Figure 4A) for RT-qPCR. If *FPN3* level in the roots is regulated by a long-distance signal from the shoots, then *FPN3* expression is expected to be higher in both sides of the split roots of plants that were initially grown in iron deficient medium, regardless of the iron conditions for each side of the split roots. Our results revealed that, regardless of the initial growth conditions, significantly more *FPN3* transcripts accumulated in the split root subjected to iron-deficiency compared to its counterpart grown in iron sufficient medium (Figure 4B). This suggests that the increased level of *FPN3* transcripts in iron deficient roots is primarily driven by the local iron conditions (Figure 4B). In the split root transferred from iron sufficient conditions to iron deficient conditions, we observed the greatest increase in *FPN3* transcript levels (Figure 4B). Meanwhile, between the split roots pre-treated under iron deficiency, only the split root subjected to an additional iron deficiency exhibited higher levels of *FPN3* over the time course tested. However, in the split-root transferred from iron sufficient to sufficient conditions, it was noted that there was a 2-fold increase in *FPN3* transcript level (Figure 4B). This observation suggests that a systemic signal from the other split root, experiencing iron deficiency, might be involved, suggesting a possible contribution of the shoot-to-root signaling. Overall, our split-root results suggest that *FPN3* expression in roots is regulated by the local iron status in the roots, but some systemic signaling is involved as well.

**Figure 4.**
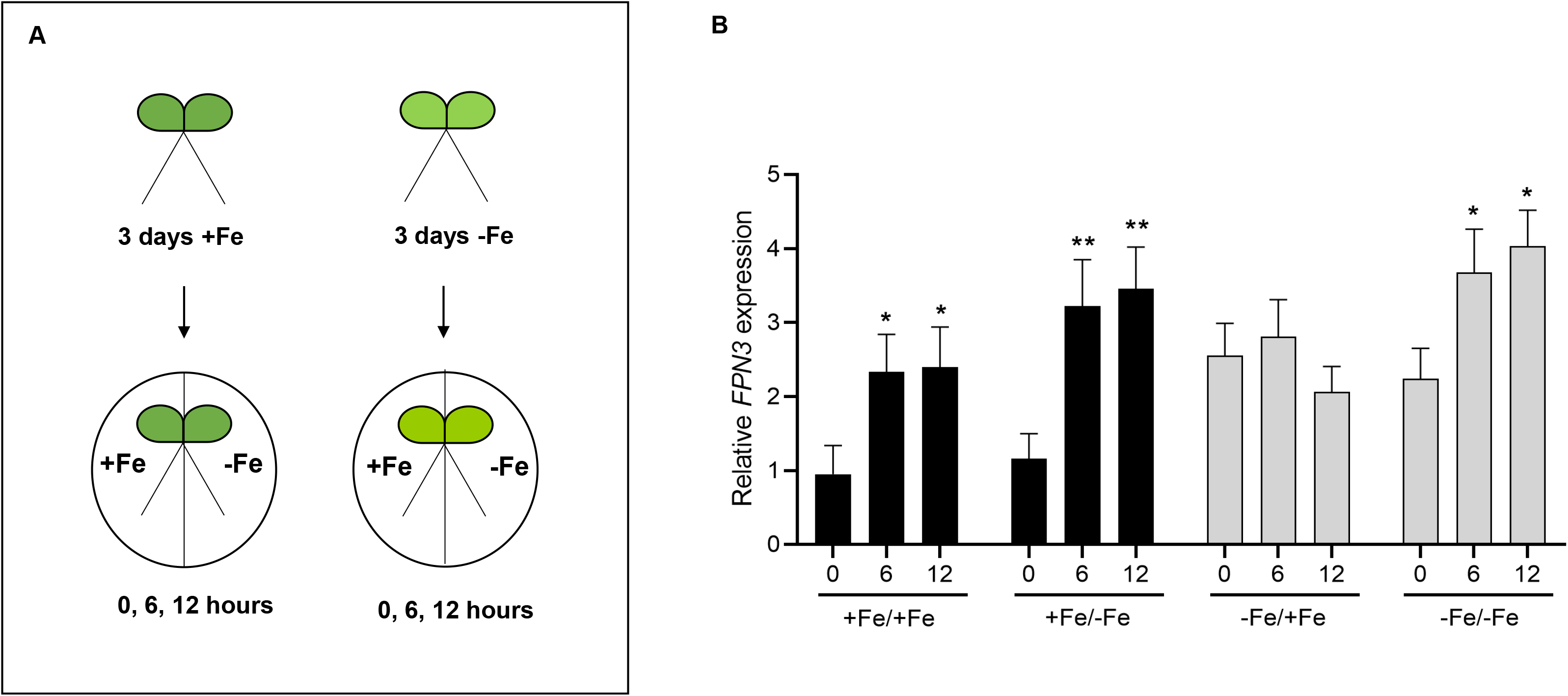
Steady state *FPN3* transcript levels in split-roots. (A) Schematic overview of iron growth conditions prior to RNA extraction. (B) *FPN3* transcript levels in split roots. Seedlings were grown under iron-deficient (−Fe) or iron-sufficient (+Fe) conditions, then each root was treated appropriately with iron-deficient or iron-sufficient media for 0, 6, or 12 hours. The iron growth conditions of each split root are denoted as: +Fe/+Fe for growth in iron sufficient conditions followed by iron sufficiency treatment; +Fe/−Fe for growth in iron sufficient conditions followed by iron deficiency treatment; −Fe/+Fe for growth in iron deficient conditions followed by iron sufficiency treatment; −Fe/−Fe for growth in iron deficient conditions followed by iron deficiency treatment. Significant differences compared to 0 hour were determined by one-way ANOVA followed by Dunnett’s test (*: p<0.05; **: p<0.01; n=5; error bars = SD).

### Growth of *fpn3* mutants is reduced under iron deficiency

We then conducted phenotypic analyses with *fpn3* plants to understand the function of FPN3 in iron homeostasis. Based on the increased *FPN3* expression in roots under iron deficiency (Figure 3I-L), we hypothesized that *fpn3* mutants might have less iron available for growth or development. To test this idea, we germinated two T-DNA insertion lines, *fpn3-1* and *fpn3-2*, which had significantly reduced levels of *FPN3* transcripts (Supplementary Figure 3) in alkaline soil, in which iron availability is drastically reduced. We observed that *fpn3* single mutants were smaller than wild type in alkaline soil, but this phenotype was recovered when watered with soluble iron (Figure 5A, 5B). As an alternative approach to verify the low iron growth phenotype, we quantified the root lengths and shoot fresh weights of the *fpn3* and wild type seedlings germinated in iron sufficient medium (Supplementary Figure 3C) or in medium without iron (Supplementary Figure 3D). Under iron deficient conditions, *fpn3* mutant seedlings exhibited decreased shoot fresh weights (Supplementary Figure 3D). The growth defect of *fpn3* on iron-deficient medium and alkaline soil consistently showed that *FPN3* is necessary for optimal growth under iron limiting conditions.

**Figure 5.**
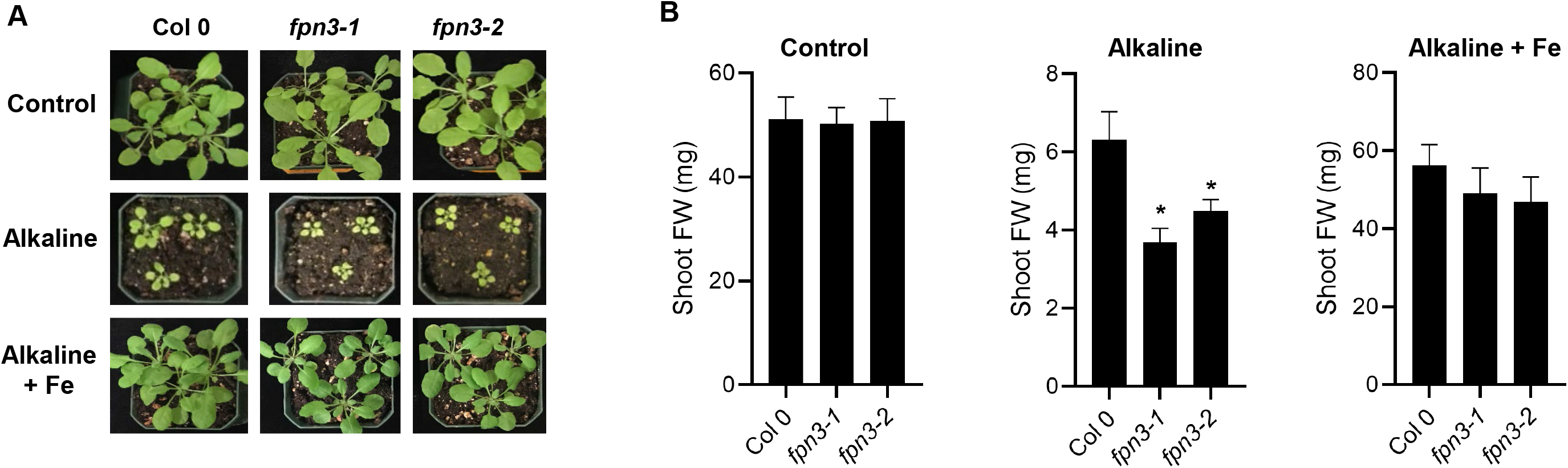
Growth of *fpn3* under iron deficient conditions. (A) Wild type (Col 0) and *fpn3* single mutants germinated and grown in control soil or alkaline soil (~pH 8) with or without iron irrigation. (B) Quantified shoot fresh weights of wild type (Col 0), *fpn3* single mutants, germinated and grown in control soil or alkaline soil. Mean values of 36 to 40 individuals are shown. Statistically significant differences compared to Col 0 are denoted (One-way ANOVA; Dunnett’s test; *: p <0.05; error bars = SD).

### Iron content is reduced in *fpn3* mutant leaves

To test if iron content of *fpn3* mutant leaves is affected, we conducted bulk elemental analysis of shoot and root tissues from plants grown under iron deficient and sufficient conditions using inductively coupled plasma mass spectrometry (ICP-MS). ICP-MS results revealed that iron content of *fpn3* mutant shoot was significantly lower than that of wild type under both iron sufficient and deficient conditions (Figure 6A). In roots, iron content was not statistically different between wild type and *fpn3* mutants under iron sufficient conditions while iron content was significantly lower in *fpn3* mutant roots than in wild type roots under iron deficiency (Figure 6B). We then performed synchrotron x-ray fluorescence imaging (SXRF) to examine the amount and distribution of iron in leaves of *fpn3* and wild type. The overall distribution of iron was not different among genotypes (Figure 6C). The elemental map of manganese and zinc were similar compared to wild type (Supplementary Figure 4). These results suggested that the function of FPN3 is necessary to properly maintain iron levels in the leaves regardless of iron conditions and in the roots under iron deficiency. We note that our ICP-MS data correlates with the expression profile of *FPN3* in roots and shoots under iron sufficient and deficient conditions (Figure 3L).

**Figure 6.**
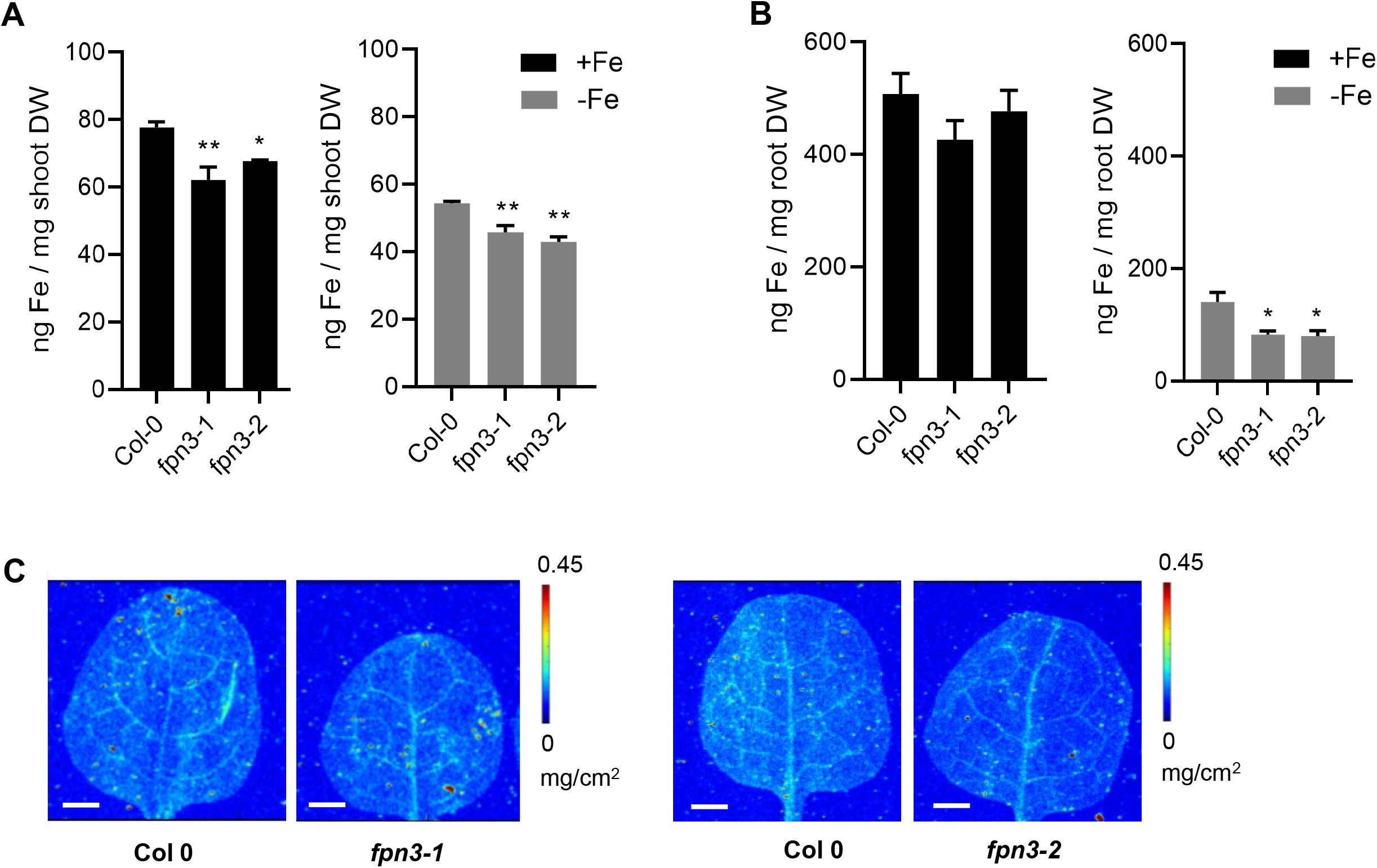
Elemental analysis of *fpn3* shoots and roots. (A) Iron content of shoot tissue from plants grown under iron sufficient (+Fe) and deficient (−Fe) conditions measured by ICP-MS. (B) Iron content of root tissue from plants grown under iron sufficient (+Fe) and deficient (−Fe) conditions measured by ICP-MS. Mean values of pooled quadruplicate ICP-MS samples are shown with error bars (SE). Significant differences compared to Col 0 are denoted (One-way ANOVA; Dunnett’s test; *: p<0.05; **: p<0.01).(C) Synchrotron x-ray fluorescence microscopy images of iron distribution in leaves of wild type (Col 0) and *fpn3* leaves. The first true leaves of 22-day old plants grown in iron sufficient conditions were imaged at a resolution of 30 × 30 μm with 0.2 sec dwell time. Representative images of leaves from three individuals are shown. Scale bars = 1 mm.

### FPN3-GFP is dual-targeted to the mitochondria and plastids in Arabidopsis

To determine the subcellular localization of FPN3, we generated transgenic Arabidopsis plants that stably express *35Sp-FPN3-GFP* and examined the subcellular localization of FPN3-GFP in roots after staining with MitoTracker Red, a mitochondrial dye. In transgenic roots, FPN3-GFP signal co-localized with the fluorescence of MitoTracker Red (Figure 7A-C). Based on quantification, 83% of the GFP signal co-localized with the mitochondrial staining. The FPN3-GFP signal that did not overlap with the mitochondrial staining (Figure 7C) suggested that FPN3-GPF might be dual-targeted. To confirm the localization of FPN3-GFP in chloroplasts, we also generated transgenic Arabidopsis plants stably expressing both *35Sp-FPN3-GFP* and *35Sp-pt-RFP,* a plastid marker with the transit peptide of rubisco small subunit from tobacco fused to RFP (Nelson *et al.*, 2007). We observed 72% co-localization of FPN3-GFP with the plastid marker (Figure 7D-F). Overall, our localization results from both stable transgenic lines confirm the plastid localization of FPN3-GFP previously reported by Conte et al. (2009) and provide further evidence that FPN3 is dual-targeted.

**Figure 7.**
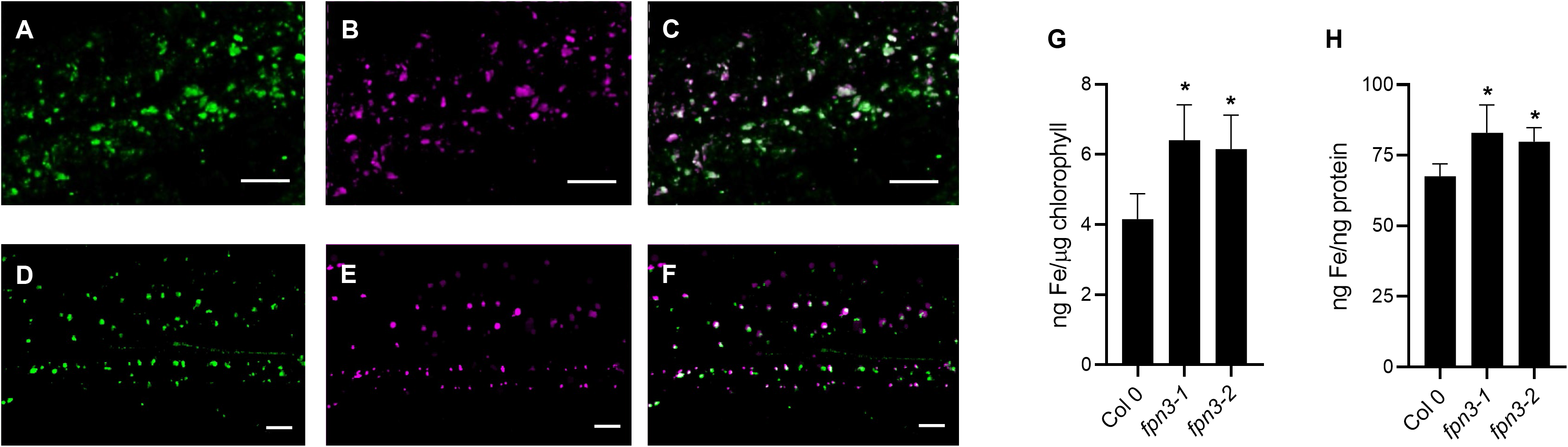
Subcellular localization of FPN3-GFP and elemental analysis of *fpn3* mitochondria and chloroplasts. (A-C) Confocal image of an optical section of Arabidopsis root stably expressing *FPN3-GFP* (A) and stained with MitoTracker Red (B). (C) Overlay image of (A) and (B) shows the co-localization of FPN3-GFP with MitoTracker Red. (D-F) Confocal image of an optical section of Arabidopsis root stably co-expressing *FPN3-GFP* (D) and *pt-RFP* (E). (F) Overlay image of (D) and (E) shows the co-localization of FPN3-GFP with *pt-RFP*. Scale bars = 25 μm. (G, H) Elemental analysis of chloroplasts and mitochondria isolated from *fpn3* shoots. (G) Iron content of chloroplasts measured by ICP-MS and normalized with chlorophyll content. (H) Iron content of mitochondria normalized with mitochondrial protein. Mean values of quadruplicate chloroplast samples and triplicate mitochondrial samples are shown with error bars (SD). Significant differences compared to the wild type, Col 0, are denoted (One-way ANOVA followed by Dunnett’s test; *: p<0.05).

### Mitochondria and chloroplasts of *fpn3* accumulate iron

Based on the subcellular localization of FPN3 in the mitochondria and chloroplasts (Figure 7A-F) and our yeast results that suggested FPN3 is exporting iron from these organelles (Figure 2), we predicted that iron would accumulate in the mitochondria and chloroplasts of *fpn3*. Thus, we isolated these organelles from *fpn3* and wild type shoots. To examine the purity of our samples, Western blots were done with the chloroplasts or mitochondrial samples using antibodies against cytosolic or organellar markers (Supplementary Figures 5, 6). We then quantified metal concentrations using ICP-MS. As predicted, chloroplasts and mitochondria from both *fpn3-1* and *fpn3-2* contained significantly more iron compared to their counterparts in wild type (Figure 7G, 7H). The accumulation of iron in *fpn3* mutant chloroplasts and mitochondria provides strong evidence that supports the role of FPN3 in exporting iron from the mitochondria and chloroplast.

### The expression of iron responsive genes is altered in the *fpn3* mutants

Trafficking of iron between subcellular compartments is crucial for cellular iron homeostasis. Because mitochondria and chloroplasts prepared from leaves of *fpn3* mutants accumulate more iron, while leaves of the mutant accumulate less iron compared to wild type, we predicted that transcript levels of organellar iron responsive genes would be affected in *fpn3* mutant lines. We first analyzed the expression of vegetative ferritin genes, *FERRITIN1 (FER1)*, *FER3*, and *FER4*, which transcripts accumulate under iron sufficiency and decrease when iron is low (Petit *et al.*, 2001; Arnaud *et al.*, 2006). We found that transcript levels of *FER1* and *FER3*, which encode plastid-localized ferritins, were significantly lower in iron-sufficient shoots of *fpn3* compared to wild type (Figure 8), which is consistent with our results showing less iron accumulation in *fpn3* shoots (Figure 6A). In contrast, the expression of *FER4,* a dual-targeted ferritin that localizes to mitochondria and plastids, was approximately 2 and 2.9-fold higher in *fpn3-1* and *fpn3-2*, respectively, compared to wild type (Figure 8).

**Figure 8.**
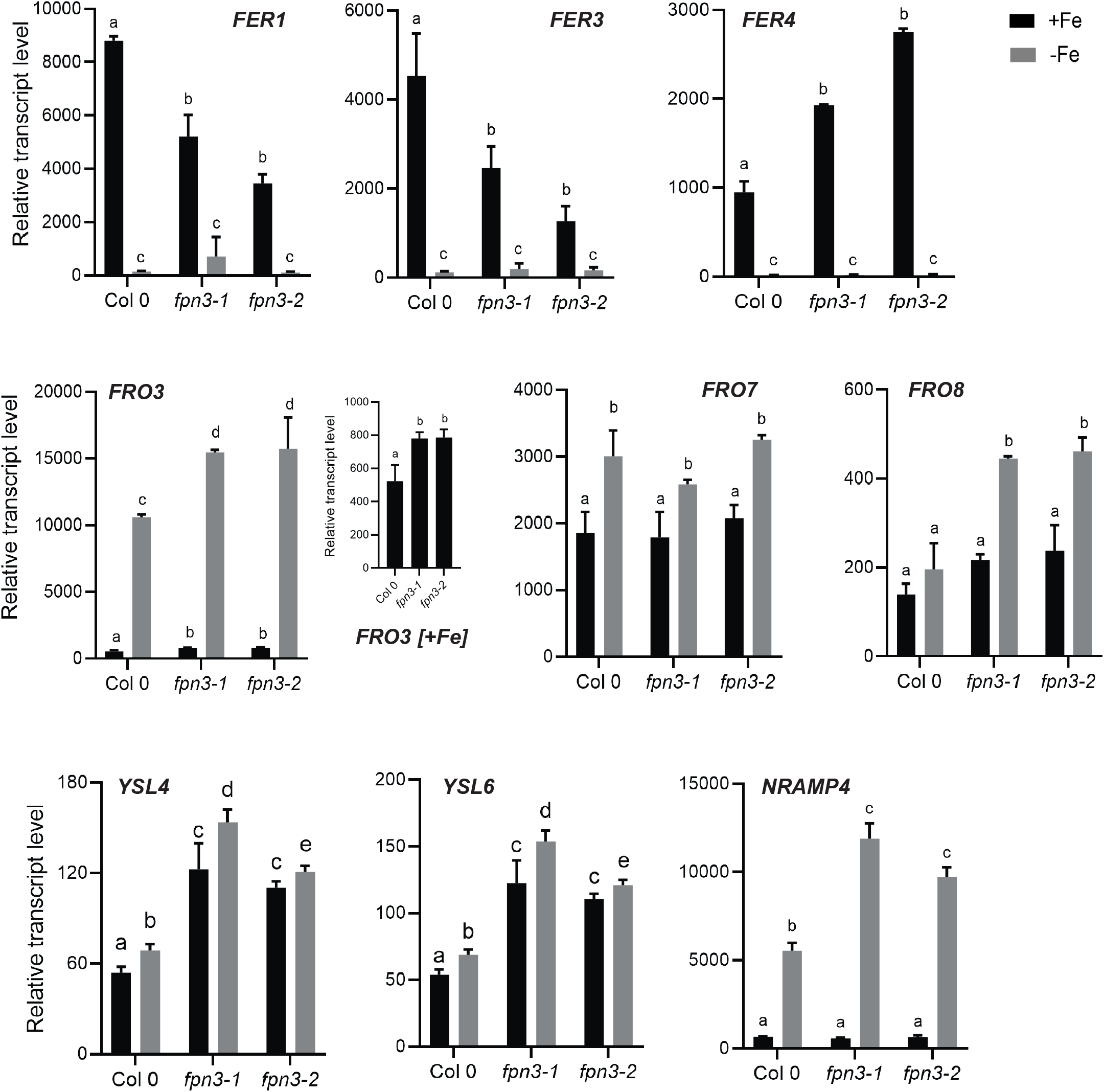
Expression of organellar iron homeostasis genes in *fpn3* shoots. Normalized steady-state level transcripts detected by NanoString. Seedlings were grown in B5 without sucrose for 2 weeks and transferred to iron-deficient or iron-sufficient media for 3 days. Mean values of normalized transcript levels are shown. Significant differences compared to Col 0 samples from the same treatment are denoted (One-way ANOVA; Dunnett’s test; *: p<0.05; n=4; Error bars = SD).

*FRO3* encodes a mitochondrial ferric chelate reductase and has been used as a marker for iron deficiency (Mukherjee *et al.*, 2006). *FRO3* expression was about 46% and 31% higher in iron-deficient shoots of *fpn3-1* and *fpn3-2,* respectively, compared to wild type (Figure 8). In addition, *FRO3* expression was significantly higher in *fpn3* shoots than in wild type under iron-sufficient conditions (Figure 8). Along with the expression profiles of *FER1* and *FER3*, elevated *FRO3* expression levels in *fpn3* shoots is consistent with less cytosolic iron (Figure 6). Expression of another mitochondrial ferric chelate reductase gene, *FRO8*, was ~30% higher in iron-deficient *fpn3* shoots than in iron-sufficient *fpn3* shoots, and was increased 2-fold in *fpn3* shoots compared to wild type shoots under iron-deficient conditions (Figure 8). In contrast to the mitochondrial ferric chelate reductase genes, expression of *FRO7*, which encodes the chloroplast ferric chelate reductase, was not significantly different between the *fpn3* mutants and wild type; *FRO7* transcript levels were increased in iron deficiency, but similar across all three genotypes (Figure 8). We noted that the expression of *YSL4* and *YSL6*, which have been reported to encode effluxors of iron-NA complexes from the chloroplasts (Divol *et al.*, 2013) or vacuoles (Jaquinod *et al.*, 2007; Conte *et al.*, 2013), was significantly increased in *fpn3* mutant shoots as compared to wild type (Figure 8). *NRAMP4* encodes a vacuolar iron effluxer and is induced by iron deficiency (Lanquar *et al.*, 2005). Compared to wild type, *NRAMP4* expression was significantly higher in iron-deficient shoots of *fpn3* (Figure 8). This observation is in accordance with the expression profiles of *FER1, FER3* and *FRO3* in shoots, which suggested that *fpn3* shoots have less iron than wild type shoots, and our elemental analysis data, which showed that *fpn3* shoots contain less iron than wild type shoots (Figure 6).

### *fpn3* mitochondria exhibit abnormal morphology under iron deficient conditions

Abnormal chloroplast ultrastructure has been observed in mutants defective in iron metabolism of multiple plants species grown under iron deficiency (Bogorad *et al.*, 1959; Duy *et al.*, 2007; Platt-Aloia *et al.*, 1983; Stocking, 1975; Vigani *et al.*, 2015). In addition to chloroplasts, the ultrastructure of mitochondria has also been shown to be affected by iron deficiency in plants (Pascal and Douce, 1993; Vigani *et al.*, 2015). Therefore, we examined the ultrastructure of chloroplasts and mitochondria in *fpn3* and wild type leaves using transmission electron microscopy (TEM). Our TEM images revealed that mitochondria of iron-deficient *fpn3* mutants were enlarged and the outer and inner membranes appeared deformed (Figure 9C), but such phenotypes were not detected under iron sufficient conditions (Figure 9A). By comparing the quantified area of mitochondria, we determined that *fpn3* mitochondria were significantly larger than wild type mitochondria under iron deficient conditions (Figure 9D), but not under iron sufficient conditions (Figure 9B). Overall, the iron-dependent mitochondrial ultrastructure phenotype (Figure 9) strongly suggests that FPN3 function is critical for mitochondria under iron deficient conditions.

**Figure 9.**
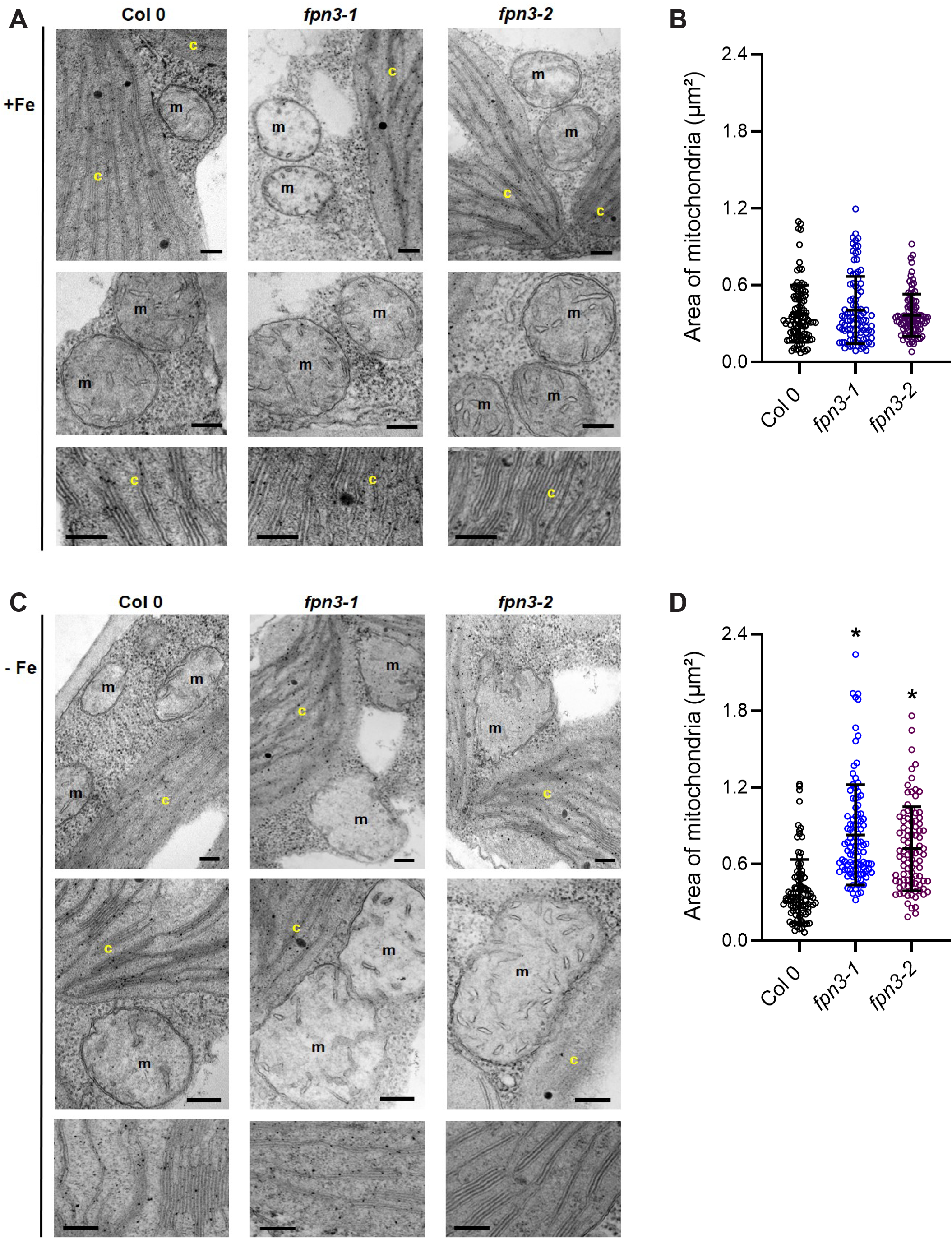
Mitochondria and chloroplast ultrastructure of *fpn3*. (A) TEM images of the wild type, Col 0, *fpn3-1*, and *fpn3-2* leaf sections with mesophyll cells from iron sufficient conditions. Close-up images of mitochondria (middle row) and thylakoids (bottom row) are shown. (B) Area of mitochondria from iron sufficient conditions. data point represents the quantified area from each mitochondrion section, and the lines represent mean values with SD. Ninety-four to 105 mitochondria from 38 to 41 cells were quantified. (C) TEM images of iron deficient wild type, *fpn3-1*, and *fpn3-2* leaf sections with mesophyll cells. Chloroplasts (c) and mitochondria (m) are labeled in the images. Scale bars = 200 nm. (D) Area of mitochondria from iron deficient conditions. Each data point represents the quantified area from each mitochondrion section, and the lines represent mean values with SD. Ninety-six to 113 mitochondria from 40 to 57 cells were quantified. (one-way ANOVA; Tukey’s multiple comparison test’ *: p<0.01).

## DISCUSSION

### FPN3 is a dual-targeted mitochondrial/chloroplast iron exporter

FPNs have been extensively studied in vertebrate species, in which the only FPN is localized to the plasma-membrane and exports iron from the cell into the plasma (Drakesmith *et al.*, 2015). In many plant species, multiple FPN/IREG paralogs are present and localize to different sub-cellular compartments. FPN1/IREG1 in *Arabidopsis thaliana* (Morrissey *et al.*, 2009) and the nodule-specific FPN2 in *Medicago truncatula* (Escudero *et al.*, 2020) are targeted to the plasma membrane, whereas the Arabidopsis FPN2/IREG2 (Morrissey *et al.*, 2009; Schaaf *et al.*, 2006), IREG1 of the nickel hyperaccumulator *Psychotria gabriellae* (Merlot *et al.*, 2014), and buckwheat IREG (Yokosho *et al.*, 2016) were identified on the tonoplast. Although MAR1 (alias FPN3/IREG3) was reported to be targeted to the chloroplasts (Conte *et al.*, 2009), our study shows that is dual-targeted to mitochondria and chloroplasts (Figure 7A-F).

This study suggests that FPN3 is exporting iron from the mitochondria and chloroplasts. FPN family members identified to date transport iron *from* the cytoplasm. As metal transporters of the same family generally retain the direction of transport in respect to the cytoplasm, an organellar FPN is likely to transport iron into the organellar lumen. However, the endosymbiotic origin of mitochondria and chloroplasts (Roger *et al.*, 2017; Zimorski *et al.*, 2014; Yoon *et al.*, 2004) suggests that FPN3 might efflux iron from its ancestral cytoplasm, i.e., from the organellar lumen into the eukaryotic cytoplasm. Under the hypothesis that FPN3 is of bacterial origin, its directionality is consistent with iron export from its ancestral cytoplasm. Based on the subcellular localization of FPN3 (Figure 7A-F) and the endosymbiotic origin of mitochondria and chloroplasts (Roger *et al.*, 2017; Zimorski *et al.*, 2014; Raven and Allen, 2003; Rockwell *et al.*, 2014), it is reasonable to speculate that FPN3 diverged from a bacterial ancestor. Prokaryotic ferroportins have not been extensively studied, but a bacterial FPN from *Bdellovibrio bacteriovorus* has been reported (Bonaccorsi di Patti *et al.*, 2015; Taniguchi *et al.*, 2015).

### The physiological role of FPN3

FPN3 was previously reported as MAR1 and proposed to opportunistically transport aminoglycoside class antibiotics (Conte *et al.*, 2009; Conte and Lloyd, 2010). In this report, we add new information that is critical to understanding the physiological role of FPN3/MAR1. First, our results show that FPN3 is dual-targeted to the mitochondria and chloroplasts and affects mitochondrial morphology under iron deficiency (Figures 7, 9). Conte et al (2009) reported that MAR1 was targeted to chloroplasts. Here, we observed that FPN3-GFP is co-localized with mitochondrial and plastid markers (Figure 7A-F). Furthermore, we detected iron accumulation in both mitochondria and chloroplasts of *fpn3* (Figure 7G, H) and observed abnormal mitochondrial morphology in iron deficient *fpn3* mutant leaves (Figure 9). Our TEM results indicate that FPN3 plays the essential role in maintaining iron homeostasis in the mitochondria. It is also possible that the chloroplasts may be better equipped with mechanisms that compensate for defective FPN3 function under iron deficient conditions. For example, given the proposed function of YSL4/6 in chloroplasts (Divol *et al.*, 2013), we speculate that YSL4/6 might be compensating for the loss of *FPN3* in plastids. Alternatively, based on their proposed role in the vacuole (Conte *et al.*, 2013), the increased YSL4/6 transcript levels in *fpn3* is consistent with the higher level of *NATURAL RESISTANCE-ASSOCIATED MACROPHAGE PROTEIN 4 (NRAMP4)* expression observed in *fpn3* (Figure 8). In addition, three ferritin genes are expressed in chloroplasts (*FER1, FER3* and *FER4*), while *FER4* is the only mitochondrial ferritin.

Second, we show that FPN3 exports iron from mitochondria and chloroplasts and most likely transports ferrous iron ions. Conte et al (2009) speculated that MAR1 may be importing the polyamine iron chelator, nicotianamine (NA), based on its structural similarity to aminoglycoside class antibiotics. However, the presence of the highly conserved iron ion binding residues in FPN3/IREG3 (Figure 1) and the iron toxicity phenotypes of yeast cells expressing *FPN3* that are analogous to yeast cells expressing *MMT1/2* (Figure 2) strongly support the idea that ferrous iron ions are FPN3/IREG3 substrates. Furthermore, our results support the role of FPN3/IREG3 as an iron exporter of mitochondria and chloroplasts, based on the yeast data (Figure 2) and iron accumulation quantified by ICP-MS with isolated chloroplasts and mitochondria (Figure 7G, H). It is noteworthy that in a recent study, BnMAR1, the FPN3/IREG3 ortholog in *Brassica napus*, was speculated to be involved in iron release from the plastids based on the correlation between its gene expression and chloroplast iron content (Pham *et al.*, 2020). Although the mechanism of transport via FPN has not been fully understood, structural and biochemical studies indicated that the bacterial FPN3 ortholog, BdFPN, transports iron and other divalent cations along their concentration gradient in a uniporter-like manner (Taniguchi *et al.*, 2015). We noted that the content of other essential metals, such as manganese, zinc, and copper, was also affected in *fpn3* chloroplasts and mitochondria (Supplementary Figure 7), which suggests that FPN3 might transport other metals in addition to iron. In particular, zinc accumulation was significantly higher in both mitochondria and chloroplasts of *fpn3* than in wild type (Supplementary Figure 7) and SXRF indicated that *fpn3* leaves contained less zinc than wild type leaves (Supplementary Figure 4). While manganese and copper accumulation in *fpn3* mitochondria and chloroplasts was also significantly different from that of wild type, this may reflect secondary effects because the pattern of changes was not consistent between the two organelles (Supplementary Figure 7). Further studies will be necessary to test if other metals could also be transported by FPN3.

Third, we show that *FPN3* is constitutively expressed regardless of the iron conditions in the shoots, whereas its transcript levels are increased by iron deficiency in the roots. We note that our data contradicts the results of Conte et al (2009), who reported the down-regulation of *MAR1* expression in low iron conditions by RT-qPCR with whole seedlings. However, multiple transcriptomics datasets corroborate our results and show that *FPN3/IREG3* transcript level is higher in wild type roots under iron deficiency (Mai *et al.*, 2016; Dinneny *et al.*, 2008; Yang *et al.*, 2010; Park *et al.*, 2019; Khan *et al.*, 2018; Buckhout *et al.*, 2009) but is not iron-regulated in shoots (Khan *et al.*, 2018; Rodríguez-Celma *et al.*, 2013). The differential expression of *FPN3* in roots and shoots under iron-deficient or iron-sufficient conditions (Figure 3G-L) implies that FPN3 might contribute to different physiological roles in roots and shoots. Genes involved in iron acquisition or mobilization are generally induced under iron deficiency (Connolly *et al.*, 2002; Varotto *et al.*, 2002; Vert *et al.*, 2002; Lanquar *et al.*, 2005; Thomine *et al.*, 2003), whereas steady-state transcript levels of multiple metal homeostasis genes that load iron into subcellular organelles remain unaffected by the availability of metals in the environment (Kim *et al.*, 2006; Zhang *et al.*, 2012; Desbrosses-Fonrouge *et al.*, 2005; Klaumann *et al.*, 2011). Our split root data revealed that the transcript level of *FPN3* in roots is regulated in response to the local availability of iron, but also indicated that systemic signals also contribute to *FPN3* regulation (Figure 4). This suggests that FPN3 is releasing iron from chloroplasts/plastids and mitochondria in the roots when iron deficiency in the rhizosphere is sensed, while responding to iron levels in the shoots. We speculate that FPN3 may be releasing iron into the root cytoplasm so that it can be reallocated within the cell to help cope with iron deficiency and to adjust metabolic needs under iron deficiency. As a response to iron deficiency, metabolic pathways in plants, including those that occur in chloroplasts and mitochondria, are coordinately remodeled in both roots and shoots (López-Millán *et al.*, 2013; Thimm *et al.*, 2001; Rodríguez-Celma *et al.*, 2013). In the shoots, *FPN3* is constitutively expressed regardless of the plant iron status (Figure 3G-L) and throughout development according to transcriptomic datasets (Winter *et al.*, 2007). FPN3 might be releasing iron to protect from iron-induced oxidative stress in the shoots. It has been proposed that plant ferritins play a major role in preventing iron-induced oxidative stress in plants (Ravet *et al.*, 2009). *FER4* encodes a dual-targeted ferritin that localizes to mitochondria and plastids. FER4 is the only mitochondrial ferritin, as mitochondria of *fer4* cells are devoid of ferritins (Tarantino, Santo, *et al.*, 2010). Thus, the pronounced increase in *FER4* transcripts is likely caused by *fpn3-1* and *fpn3-2* mutant plants accumulating more iron in the mitochondria and chloroplasts and, in response, higher level of *FER4* transcripts to prevent iron-induced oxidative damage.

Although *FPN3* does not appear to be iron-regulated at the transcriptional level in the shoots, the abnormal mitochondria morphology detected in iron deficient *fpn3* leaves suggest a critical role for FPN3 in the mitochondria under iron deficiency (Figure 9). Because *FRO8* is not regulated by the iron status of wild type plants (Figure 8;(Mukherjee *et al.*, 2006), the higher level of *FRO8* in iron-deficient *fpn3* shoots (Figure 8) may also imply a dysregulation of iron-dependent mitochondrial function. While the mechanism behind the enlarged mitochondria of iron deficient *fpn3* shoots remains to be understood, it is noteworthy that impaired ion homeostasis is known to affect mitochondrial volume (Szabo and Zoratti, 2014; Teardo *et al.*, 2015). Swollen mitochondria have been observed upon anaerobic treatment, decreased metabolism, or oxidative stress (Vartapetian *et al.*, 2003; Yoshinaga *et al.*, 2005; Lee *et al.*, 2002). We postulate that oxidative stress or metabolic constraints caused by dysregulated iron homeostasis in iron deficient *fpn3* mitochondria might have resulted in the compromised mitochondrial ultrastructure. We further speculate that the reduced copper level in *fpn3* mitochondria (Supplementary Figure 7B) may indicate that *fpn3* mitochondria are more susceptible to oxidative stress than *fpn3* chloroplasts. A recent study has shown that mitochondria accumulates oxygen under low copper due to impaired cytochrome c function (Ishka and Vatamaniuk, 2020).

### Implications of a mitochondrial iron exporter

Mitochondrial metabolism was found to be less affected than chloroplast metabolism under mild iron deficiency (Hantzis *et al.*, 2017). However, it is noteworthy that improper FPN3 function appears to have a greater impact on the mitochondria than chloroplasts (Figure 9). The mitochondrial phenotype of *fpn3* signifies the importance of iron export from the mitochondria under iron deficient conditions. It has been well-established that chloroplasts are the most iron-rich organelle in plant cells and vacuoles serve as the major iron storage compartment, and several studies have provided evidence that mitochondria can contribute as an additional intracellular iron reservoir. In yeast *Δccc1* mutant cells that lack the vacuolar iron importer Ccc1, overexpressing *MRS3* and *MRS4*, which encode mitochondrial iron importers of the mitoferrin family, suppressed iron toxicity (Lin *et al.*, 2011). Furthermore, MMT1 and MMT2, which belong to the cation diffusion facilitator (CDF) family, export mitochondrial iron and overexpression of *MMT1/2* results in multiple phenotypes that support the idea that mitochondria could function as an iron storage organelle (Li *et al.*, 2014). It is interesting to note that mitochondrial iron import mediated by mitoferrins is common across multiple organisms, such as in yeast, mammals, worm, fruit fly, rice, and Arabidopsis (Paradkar *et al.*, 2009; Shaw *et al.*, 2006; Metzendorf *et al.*, 2009; Ren *et al.*, 2012; Bashir *et al.*, 2011; Mühlenhoff *et al.*, 2003; Jain *et al.*, 2019). Unlike mitochondrial iron import, we speculate that mechanisms to export mitochondrial iron are more diverse. FPNs do not exist in yeast and mitochondrial FPNs have not been found in vertebrates or invertebrates, to our knowledge. In mammalian mitochondria, an ATP-binding cassette (ABC) transporter, ABCB8, is involved in exporting iron (Ichikawa *et al.*, 2012). The presence of a mitochondrial iron exporter is necessary, but not sufficient to suggest that mitochondria might function as a subcellular iron store. Nevertheless, our work shows that FPN3 is regulating mitochondrial iron and implies a potential role of mitochondria as an iron reservoir. It is possible that *fpn3* mutants might be increasing FER4 to protect from iron toxicity due to the lack of an iron exporter. Furthermore, the higher expression of *FER4* in *fpn3* than in wild type (Figure 8) is also consistent with the view that mitochondria could store iron. We postulate that mitochondrial ferritin may play a more dynamic function to assist iron storage in addition to protecting mitochondria from iron-induced oxidative damage.

In conclusion, our work has advanced our understanding of the physiological roles of FPN3 by providing evidence that FPN3 is an iron exporter dual-targeted to mitochondria and chloroplasts. Our TEM results imply that FPN3 function is more crucial for mitochondria, but further studies are necessary to elucidate the mechanisms underlying the phenotypes observed and the potential effect of FPN3 on mitochondrial and chloroplast metabolism and oxidative stress. Future work should also address potential crosstalk among chloroplasts, mitochondria, and vacuoles to comprehensively understand organellar iron trafficking and homeostasis in plant cells.

## MATERIALS AND METHODS

### Yeast high iron growth assays

For growth tests in high iron media, *pRS315-FPN3*, *pRS315-MMT1/2*, or *pRS315* vector alone was transformed into *Δccc1Δmmt1/2* cells. For spot assays, serial dilutions of overnight yeast cultures were spotted onto SD-Leu plates with appropriate auxotrophic supplements containing 2 mM iron (2 mM ferrous ammonium sulfate hexahydrate). The negative control plates contained media with no added iron. Cells growth at 30°C was monitored over 2-5 days. For growth assays in liquid culture, overnight yeast cultures were inoculated to a starting OD_600_ of 0.05 into SD-Leu with or without 2 mM iron supplementation and grown in a shaker incubator at 30°C, and cell density was measured at OD_600_ at the indicated time points.

### Aconitase assay

Wild type DY150 cells were transformed with *FPN3*, *MMT1/2*, or *pRS315* vector alone. These transformants were inoculated in appropriate selective media (SD -Leu), grown overnight at 30°C, and sub-cultured in SD-Leu supplemented with 250 μM FeSO_4_ until the cells reached mid-log phase. The subcultures were harvested and washed with cold phosphate-buffered saline (PBS), and protein lysate was prepared using PBS with protease inhibitor (Pierce) via homogenization with glass beads for 10 minutes at 4°C. Samples used in the aconitase assay were normalized with total protein concentration. Aconitase activity was assayed and quantified using the BioAssay Systems EnzyChrom Aconitase Assay Kit, following manufacturer’s instructions.

### c-GDO assay

Wild type DY150 cells were co-transformed with *FPN3, MMT1/*2, or *pRS315* and the *c-GDO* plasmid, and grown as described above for the aconitase assay. Cell lysates were prepared with these transformants by glass bead homogenization, and GDO activity was assayed as described by Li et al (2014). The assay reaction consisted of 20 mM Tris-HCl (pH 8.0) and 0.1 mM 2,3-dihydroxy-benzoic acid (gentisic acid) as the substrate. The enzyme activity was determined based on the absorbance measured at 340 nm using an extinction coefficient of 10.2 cm^−1^mm^−1^.

### Plant materials and growth conditions

Two independent alleles of *fpn3* T-DNA insertion lines, *fpn3-1* (SALK_034189) and *fpn3-2* (SALK_009286), were obtained from the Salk collection (Alonso *et al.*, 2003). RT-qPCR verified that *FPN3* expression was drastically reduced in these lines (Supplementary Figure 3). Arabidopsis plants were grown at 22 °C under a 16/8-hour light/dark cycle in the Conviron A1000 growth chamber.

### Plant growth test under iron-deficient conditions

For alkaline soil tests, seedlings were germinated and grown in normal soil (pH ~5.8), alkaline soil (pH ~8), or alkaline soil watered with soluble iron (0.5 mM Fe-EDDHA). Alkaline soil was prepared by adding approximately 1.3 g of calcium oxide per liter of soil solution (Kim *et al.*, 2006). For growth tests on plates, seedlings were germinated vertically for 12 days on either iron-deficient media (Murashige and Skoog (MS) without iron (Caisson MSP33) supplemented with 300 μM ferrozine) or iron-sufficient media (MS with 100 μM FeNa-EDTA (Caisson MSP34). Seedlings were photographed, and ImageJ was used to quantify root lengths. Shoots were separated and weighed on the MT5 Analytical Microbalance (Mettler Toledo).

### Generation of *FPN3p-GUS* transgenic lines and histochemical staining

A fragment containing 890 bp of upstream sequence and the *FPN3* start codon was amplified from a BAC clone, F2P16 (GenBank AF007270), obtained from the Arabidopsis Biological Resource Center (ABRC) using primers 5’-cacccactttctcttttgttagattctagttg-3’ and 5’-cattctataaat-tgattctcctcttctcc-3’ and cloned into pENTR/D-TOPO (Invitrogen). The construct was cloned into pGWB533 (Nakagawa *et al.*, 2007) using Gateway LR Clonase II (Invitrogen). The resulting *FPN3p-GUS* construct was moved into *Agrobacterium tumefaciens* strain LBA4404 and transformed into wild type Col 0 plants using the floral dip method (Clough and Bent, 1998). pGWB533 was from Tsuyoshi Nakagawa (Addgene plasmid #74872; RRID: Addgene_74872). For histochemical staining, seedlings or tissues of *FPN3p-GUS* plants were fixed and incubated with 5-bromo-4-chloro-3-indolyl β–D-glucuronide as described by (Jefferson *et al.*, 1987). At least five seedlings from three independent T2 or T3 lines were examined, and images were obtained using the LeicaM165 FC dissecting scope and Leica LAS EZ 3.4.0 software.

### Split root assays

For split-root samples for RT-qPCR, plants were prepared following the protocol described by Kumar et al. (2017) with slight modifications. Wild type plants were grown for 5 days on iron-sufficient medium, cut at the roots and grown for 7 days to induce split roots, and transferred to iron-deficient or iron-sufficient medium for 3 days. The plants were then washed with 0.1M citrate buffer and moved to split plates, in which one of the split roots was placed on iron-deficient medium and the other root on iron-sufficient medium. Plants were kept on split plates for 0, 6 and 12 hours before harvesting roots for RNA preparation.

### Generation of *FPN3-GFP* transgenic lines

The coding region of FPN3 was cloned from a BAC clone, F2P16 (GenBank AF007270), obtained from the Arabidopsis Biological Resource Center (ABRC) using primers 5’-caccatggttgtttcaatggctttgg-3’ and 5’-atttgagagagggtcgaaggag-3’ and cloned into pENTR/D-TOPO (Invitrogen). The final construct, *35Sp-FPN3-GFP*, was cloned into pGWB505 (Nakagawa *et al.*, 2007) using Gateway LR Clonase II (Invitrogen). *35Sp-FPN3-GFP* and *35Sp-pt-RFP* were co-transformed into *Agrobacterium tumefaciens* strain LBA4404, and then into wild type Col 0 using the floral dip method (Clough and Bent, 1998). pGWB505 was a gift from Tsuyoshi Nakagawa (Addgene plasmid #74847; RRID: Addgene_74847) and *35Sp-pt-rb* (Nelson *et al.*, 2007) was obtained from ABRC (stock #CD3-1000).

### Confocal microscopy

*Arabidopsis* roots stably expressing *35Sp-FPN3-GFP* were imaged with the Nikon A1 Spectral Detector Confocal with FLIM Module at the Light Microscopy Core, Institute for Applied Life Science at University of Massachusetts, Amherst. For mitochondrial co-localization roots were stained with MitoTracker Red FM (Invitrogen) t2o8label the mitochondria prior to imaging, by incubating roots in a final concentration of 1 μM MitoTracker Red FM for 30 minutes. Z-stack images were taken with FITC and TRITC channels. Images were collected and processed with NIS-Elements software.

### Chloroplast isolation

Chloroplast isolation was conducted following (Smith *et al.*, 2003). Arabidopsis plants were grown for 4 weeks on soil. Roughly 10 g of shoot tissue was harvested, rinsed in digestion buffer (20 mM MES-KOH, pH 5.2, 400 mM sorbitol, 0.5 mM CaCl_2_), and incubated in digestion enzyme solution (10 ml digestion buffer, 0.04 g Macerozyme R-10 (bioWORLD), 0.2 g Cellulase (bioWORLD) for 3.5 h under light. During digestion, a 40%:85% AT Percoll step gradient was prepared on ice. Upon completion of digestion, the solution was filtered through cheesecloth to harvest protoplasts. To isolate protoplasts, the filtered solution was suspended in digestion buffer and centrifuged at 100 *g* for 5 min at 4°C, the supernatant was discarded, then the isolated protoplasts were resuspended in protoplast resuspension buffer (20 mM MES-KOH, pH 6.0, 400 mM sorbitol, 0.5 mM CaCl_2_) before centrifuging at 100 *g* for 2 min at 4°C and removing supernatant. Next, protoplasts were resuspended in protoplast breakage buffer (20 mM tricine-KOH, pH 8.4, 300 mM sorbitol, 5 mM EDTA, 5 mM EGTA, 10 mM NaHCO_3_, 0.05 g BSA added per 50 mL of solution before use) and immediately filtered through a protoplast rupturing device (a syringe tube with the end cut off, with a 20 μm and 10 μm mesh, respectively, attached via electrical tape). Next, the broken chloroplasts were layered onto the 40%:85% AT Percoll step gradient and centrifuged at 2500 *g* for 10 min at 4°C, with the brake off. A green band was visualized at the 40%85% AT Percoll interface and harvested. The resulting solution was diluted in HEPES-sorbitol buffer, pH 8.0 (50 mM HEPES-KOH, pH 8.0, 330 mM sorbitol) and centrifuged at 700 *g* for 5 min at 4°C. After removing the supernatant, resulting chloroplasts were resuspended in HEPES-sorbitol buffer, pH 8.0. Chlorophyll content (chlorophyll *a* and *b*) was quantified with the NanoDrop One Spectrophotometer (ThermoFisher) following manufacturer’s instructions. The following equations were used to quantify chlorophyll *a* and chlorophyll *b* content, respectively: *C*_a_ = 14.85*A*^666^ – 5.14*A*^650^; *C*_b_ = 25.48*A*^650^ – 7.36*A*^666^ (Porra *et al.*, 1989; Barnes *et al.*, 1992). Purity was confirmed via Western blots with organellar markers.

### Mitochondrial isolation

Mitochondria were isolated from four-week-old Arabidopsis leaves using a protocol adapted from (Keech *et al.*, 2005). Prior to the isolation, 30mL of continuous 50% Percoll gradient (50% (v/v) Percoll, 0.3M sucrose, 10 mM TES, 1 mM EDTA, 10 mM KH_2_PO_4_, 1 mM glycine, pH adjusted to 7.5 with KOH) was centrifuged at 39000 *g* for 40 min and kept at 4°C. Roughly 40-50g of tissue was homogenized in pre-chilled grinding buffer B (0.3M sucrose, 60 m TES, 2 mM EDTA, 10mM KH_2_PO_4_, 25 mM tetrasodium pyrophosphate, 1 mM glycine, 1% (w/v) polyvinyl-pyrrolidone-40, 1% (w/v) defatted bovine serum albumin (BSA), with 50 mM sodium ascorbate and 20 mM cysteine added and readjustment of the pH to 8.0 with KOH just prior to grinding) using a hand blender, filtered through a 20-μm nylon mesh, and centrifuged at 2500 *g* for 5 minutes to remove intact chloroplasts and thylakoid membranes. The supernatant was centrifuged at 15000 *g* for 15 minutes. The pellet obtained was resuspended in wash buffer B (0.3M sucrose, 10mM TES, 10mM KH_2_PO_4_, pH adjusted to 8.0 with KOH) and gently homogenized in a chilled glass homogenizer. The resuspended pellet was then layered on top of the 50% Percoll gradient and centrifuged at 15000 *g* for 20 minutes; each 30 mL density gradient has a maximum load of mitochondria from 30g of fresh leaf tissue. After centrifugation, the mitochondria formed a whitish band near the bottom of the tube. This band was aspirated and resuspended in 0.2-0.3 mL of wash buffer B to obtain a roughly 20-fold dilution. All procedures were conducted in the cold room or at 4°C. Mitochondrial protein was quantified using DC Protein Assay (Bio-Rad).

### SXRF Imaging

The spatial distribution of iron, manganese, and zinc in hydrated leaf tissues was imaged via SXRF at the F3 station, a bending magnet beamline with multilayer monochromator, at the Cornell High Energy Synchrotron Source. Seeds were directly germinated on soil and grown at 22°C, 14 h light/10h dark photoperiod at a photo flux density of 110 μmol/m^2^/sec. The nutrient solution contained the following components: 1.25 mM KNO_3_, 0.625 mM KH_2_PO4, 0.5 mM MgSO_4_, 0.5 mM Ca(NO_3_)_2_, 10 μM Fe(III)HBED, 17.5 μM H_3_BO_3_, 3.5 μM MnCl_2_, 0.125 μM CuSO_4_, 0.25 μM ZnSO_4_, 0.05 μM Na_2_MoO_4_, 2.5 μM NaCl, and 0.0025 μM CoCl_2_. The hydroponic solution was changed every week. Data were analyzed and processed as described in Yan et al 2017. Briefly, the first true leaves from 22 day-old plants were detached immediately prior to imaging, placed in the wet chamber made between two layers of metal-free Kapton™ film and mounted onto 35 mm slide mounts. The 2D raster maps were acquired using a focused, monochromatic incident x-ray beam at 12.8 keV and photon flux of approximately 3×10^10^ photons/sec. Samples were scanned at the resolution of 30 × 30 μm and acquisition time of 0.2 sec per data point. These settings did not cause damage to plant tissues within 6-8h scans required for analysis of the full set of genotypes. Element-specific x-ray fluorescence was detected using a Vortex ME-4 Silicon Drift Detector (SDD). Calibration of XRF equipment were done using a uniform thin metal film standard during each experiment. Data were processed with Praxes, a software that employs PyMca libraries in batch mode (Solé *et al.*, 2007).

### ICP-MS analysis

Shoot iron content of Col 0, *fpn3-1,* and *fpn3-2* was measured by ICP-MS and normalized with dry weight. Plants were grown following the same procedure as the growth test on low iron plates. Dried shoot, root, chloroplast, and mitochondrial samples of Col 0, *fpn3-1,* and *fpn3-2* were digested with nitric acid and analyzed with the Perkin-Elmer NexION 350D ICP-MS at the Mass Spectrometry Core, Institute for Applied Life Science at University of Massachusetts, Amherst. Metal content was recorded in parts per billion (ppb) and normalized to the dry mass (mg; shoot and root tissue), chlorophyll content (μg; isolated chloroplast) or total protein (μg; isolated mitochondria) of each sample.

### Gene expression analyses

Total RNA isolated with the Agilent Plant RNA Minikit from root or shoot tissue of plants grown under iron-deficient or iron-sufficient conditions were used for gene expression analyses. The quantity and purity of RNA were examined using NanoDrop One (Thermo Scientific), and the integrity of RNA was assessed by electrophoresis using a bleach gel (Aranda *et al.*, 2012). For RT-qPCR, first strand cDNA was synthesized from total RNA using Quanta qScript, and qPCR was conducted using Bio-Rad SYBR Green Supermix in CFX-Connect Real-Time PCR Detection System (Bio-Rad). The relative transcript level of genes was calculated following the ΔCt method (Schmittgen and Livak, 2008), using *ACTIN2* (*ACT2*) as an internal control. Primers were designed using QuantPrime (Arvidsson et al., 2008) and the sequences are listed in Supplementary Table 1. For Nanostring, a custom CodeSet that included probes specific to iron homeostasis genes of interest was designed by NanoString. Gene expression analysis was conducted with the NanoString MAX/FLEX nCounter at Dartmouth College and nCounter SPRINT Profiler System at the Molecular Genetics Core Facility at Boston Children’s Hospital following the manufacturer’s instructions and processed using nSolver 4.0. The custom CodeSet sequences are listed in Supplementary Table 2.

### Transmission electron microscopy (TEM)

Excised cotyledons of Arabidopsis seedlings treated under iron sufficient or deficient conditions were fixed by immersing them in 2.5% glutaraldehyde/ 2% paraformaldehyde in 0.1 M Na Cacodylate buffer (pH 7.2) and incubating them under vacuum for 30 min. After this primary fixation, the samples were rinsed three times in fresh fixation buffer for 10 min. each time and were secondarily fixed with 1.0% osmium tetroxide in ddH2O (w/v) for 1hr at room temperature. The samples were then rinsed again three times in ddH2O and then placed in 1% aqueous uranyl acetate (w/v) in the refrigerator overnight (tertiary fixation). After three more rinses in ddH2O, the samples were then dehydrated through a graded series of ethanol (10%, 30%, 50%, 75%, 85%, 95%, and 100% for 3 changes). Infiltration was accomplished by running the samples through ethanol 100%: Spurr’s resin-hard formulation (75:25 / V:V) for 1 h at room temperature, then to ethanol 100%: Spurr’s resin-hard formulation (50:50 / V:V) and finally to ethanol 100%: Spurr’s resin-hard formulation (25:75 / V:V) overnight. The following day the samples were transferred through 5 changes of 100% Spurr’s epoxy resin each 1 hr long, and then placed in molds and polymerized for two days at 68° C. Ultrathin sections (approx. 70 nm thick) were collected onto 200 mesh copper support grids and contrasted with Uranyl Acetate and Lead Citrate and then imaged using a CM 10 transmission electron microscope, under 100Kv accelerating voltage. Images were recorded with a Gatan Erlangshen CCD Digital camera.

## ACKNOWLEDGEMENTS

We are grateful to Mary Lou Guerinot and Jerry Kaplan for feedback on this manuscript. We also thank Jerry Kaplan for the yeast strains and plasmids used in this study, Liangtao Li for the yeast Western blot and help with aconitase assays, Elsbeth Walker for advice on the split root experiments, Inhwan Hwang for the pCamV3-GFP and F1-ATPase-RFP plasmids, James Chambers for assistance with confocal microscopy, Britney Privett for assistance with NanoString, Steve Eyles for help with ICP-MS, and Lara Strittmatter and UMass Medical School Electron Microscopy Facility for assistance with TEM. We also thank Veronica Voronina, Joye Yang, and Aleks Merkovich for technical assistance at various stages of this project.

This work was supported by the National Science Foundation (NSF) grant to JJ (IOS-1754969) and OKV (IOS-1656321 and IOS-1754966), the Gregory S. Call Undergraduate Research Program (LK, KT, EYP, FH, JG, MC, AK, DC, CD, EMP), Doelling Undergraduate Research Fund (LK, KT, EYP, JG, AK, DC, CD, EMP), Schupf Scholars Program (JZ), and the Sarles Fellowship (MC). The transmission electron microscopy was supported by the National Center for Research Resources award (SI0OD021580) to the UMass Medical School Electron Microscopy Facility. The authors are solely responsible for the content of this paper and do not necessarily represent the official views of the National Center for Research Resources, National Institutes of Health, or NSF. CHESS is supported by the NSF and NIH/NIGMS via NSF Award DMR-1332208. The authors declare no conflict of interests.

