## Supplementary Figures for "Ferroportin 3 is a dual-targeted mitochondrial/chloroplast iron exporter necessary for iron homeostasis in Arabidopsis"

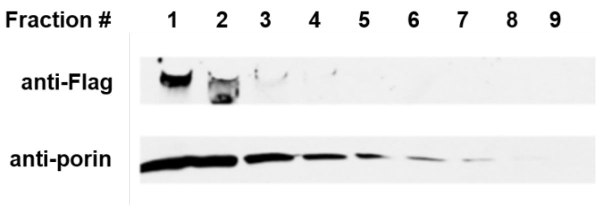


**Supplementary Figure 1.** FPN3-FLAG is targeted to the mitochondria in yeast.

Western blots with fractions containing FPN3-FLAG and the mitochondrial marker, porin. FPN3-FLAG is detected in the fractions enriched with porin.

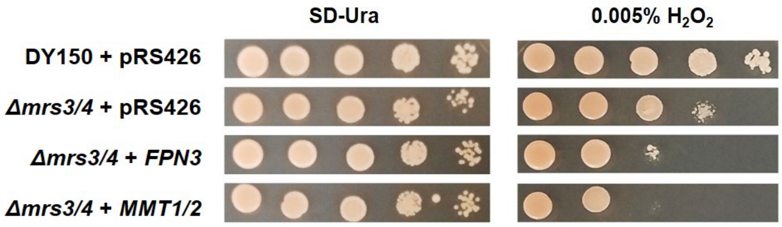


**Supplementary Figure 2. Heterologous expression of *FPN3* in *Δmrs3/4* yeast.** (A) Spot assays under oxidative stress conditions with *Δmrs3/4* cells expressing *FPN3* or *MMT1/2*. Wild type (DY150) and *Δmrs3/4* cells transformed with an empty vector, *pRS426*, were used as controls.

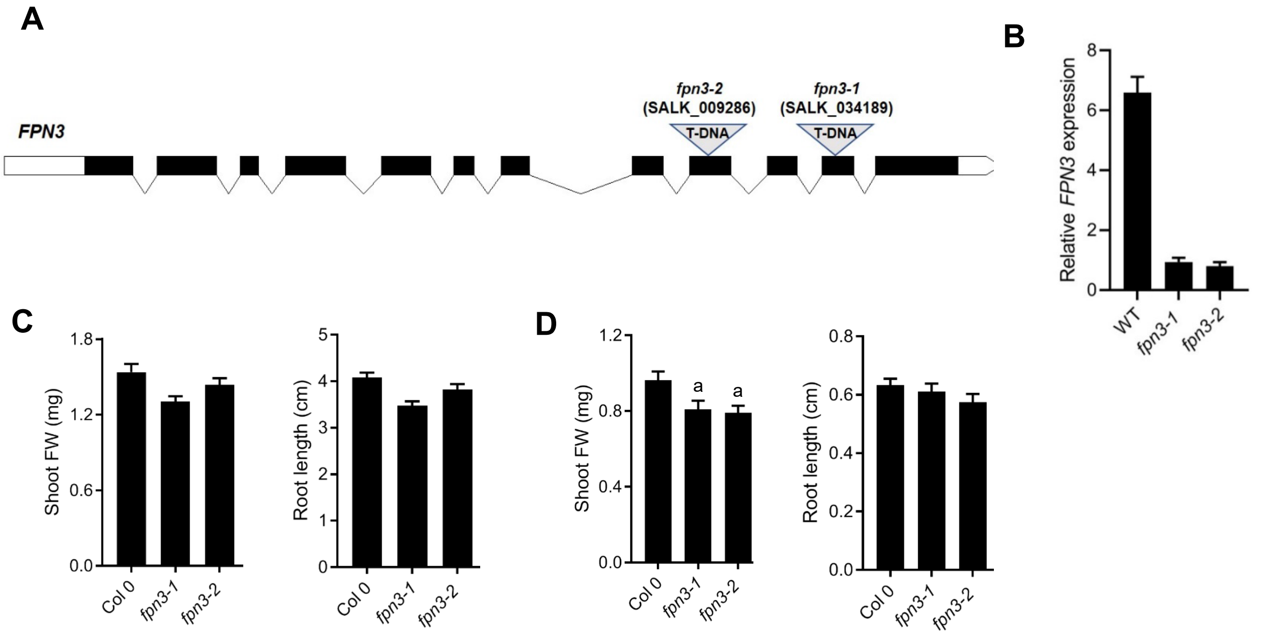


**Supplementary Figure 3.** ***FPN3*** **T-DNA insertion lines and steady-state expression level of** ***FPN3*** **in** ***fpn3*** **and** ***vit1*** **mutant lines.** (A) Schematic diagram of T-DNA insertions in *fpn3-1* (SALK_034189) and *fpn3-2* (SALK_009286). Black blocks with numbers represent exons and the gray lines are the introns. The light gray boxes before the first exon and after the last exon represent UTRs. (B) Steady state transcript level of *FPN3* in *fpn3-1* and *fpn3-2* shoots detected by RT-qPCR. Mean values of relative expression levels normalized with *ACT2* are shown (n=3; error bars=SE). The following primers were used: *FPN3* forward, 5'- GTGGGTTCTTTGCCAA-CCATGAC-3'; *FPN3* reverse, 5'-TTAGGACGGTCCAGAACTCCAG-3'; *ACT2* forward, 5'-CCAAGCTGTTCTCTCCTTGTACGC-3'; *ACT2* reverse, 5'-TCACCAGAATCCAGC-ACAAT-ACC-3. (C, D) Quantified shoot fresh weights and root lengths of wild type (Col 0), *fpn3* single mutants, germinated and grown on plates with iron-sufficient (C; 100 µM Fe) medium or iron-deficient (D; no added iron with 0.3 mM ferrozine) medium. Mean values of at least 19 individuals are shown with standard error. Statistically significant differences compared to Col 0 are denoted (One-way ANOVA; Dunnett's test; *: $p < 0.05$).

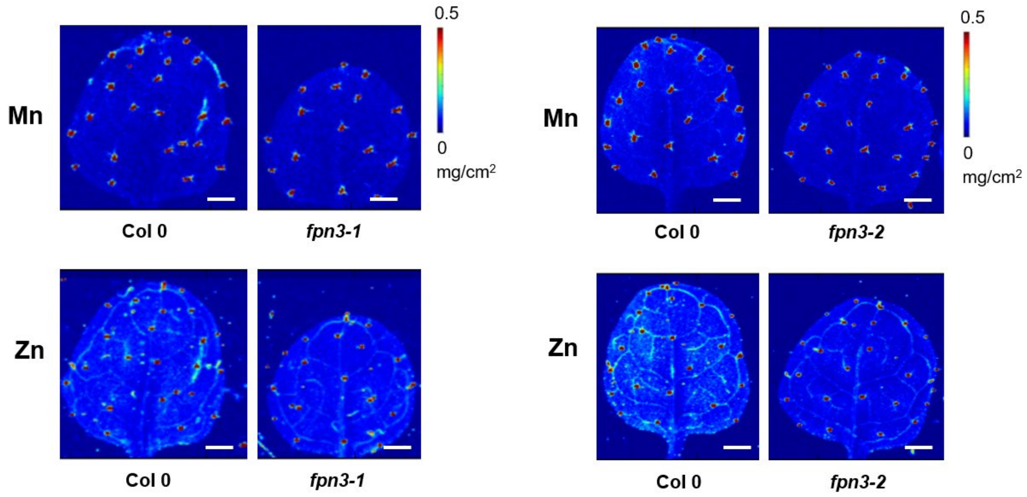


**Supplementary Figure 4. Elemental map of manganese and zinc in *fpn3-1* and *fpn3-2* leaves.**

Syncrhotron x-ray fluorescence microscopy images of the elemental distribution of manganese and zinc in leaves of wild type (Col 0) and *fpn3* single mutants. Lower resolution (40 x 40 µm; 0.2 sec dwell time) images in the first true leaves of 22 day old plants. Representative images of leaves from three individuals are shown. Scale bars = 1 mm.

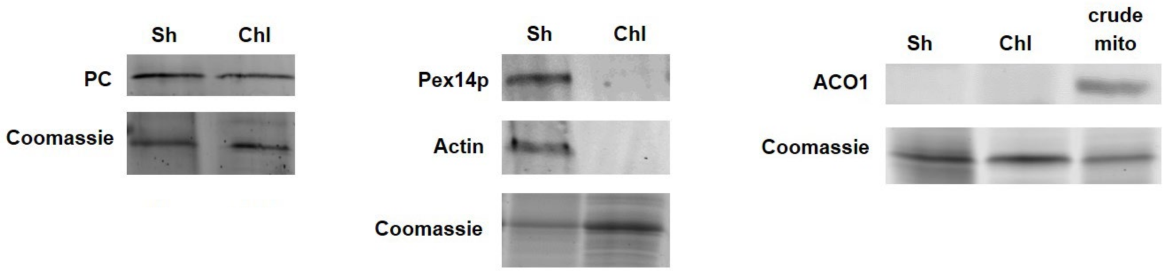


**Supplementary Figure 5. Western blots with wild type and *fpn3* chloroplast samples.**

The purity of chloroplast samples was assessed by Western blots using antibodies against the following subcellular compartment markers: plastocyanin (PC), a chloroplast marker; Pex14p, a peroxisome marker; actin, a cytosolic marker; and aconitase 1 (ACO1), a mitochondrial marker. Shoot protein (Sh) was loaded as a control alongside the chloroplast (Chl) samples. As a loading control, Coomassie staining was conducted with each set of Western blots. Because ACO1 was not detectable in our shoot samples due to its low abundance, a sample with crude mitochondrial prep (crude mito) was included as a positive control for the Western blot with anti-ACO1 antibodies.

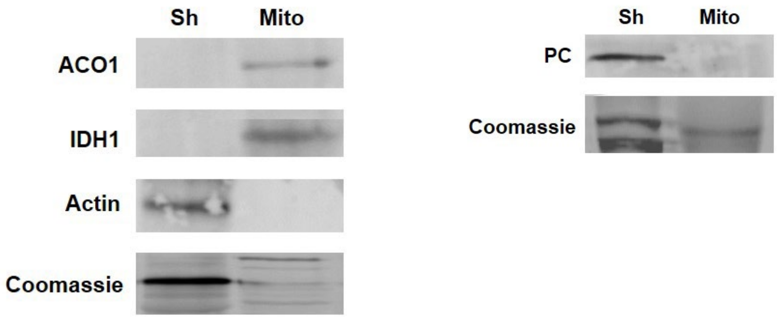


**Supplementary Figure 6. Western blots with wild type and *fpn3* mitochondria samples.**

The purity of mitochondrial samples was assessed by Western blots using antibodies against the following subcellular compartment markers: aconitase 1 (ACO1) and isocitrate dehydrogenase (IDH), mitochondrial markers, plastocyanin (PC), a chloroplast marker; Pex14p, a peroxisome marker; and actin, a cytosolic marker. Shoot protein (Sh) was loaded as a control alongside the chloroplast (Chl) samples. As a loading control, Coomassie staining was conducted with each set of Western blots. Because a small amount of ACO1 may also be found in the cytoplasm, antibodies against an additional mitochondrial marker, IDH, was also used to confirm our mitochondrial sample.

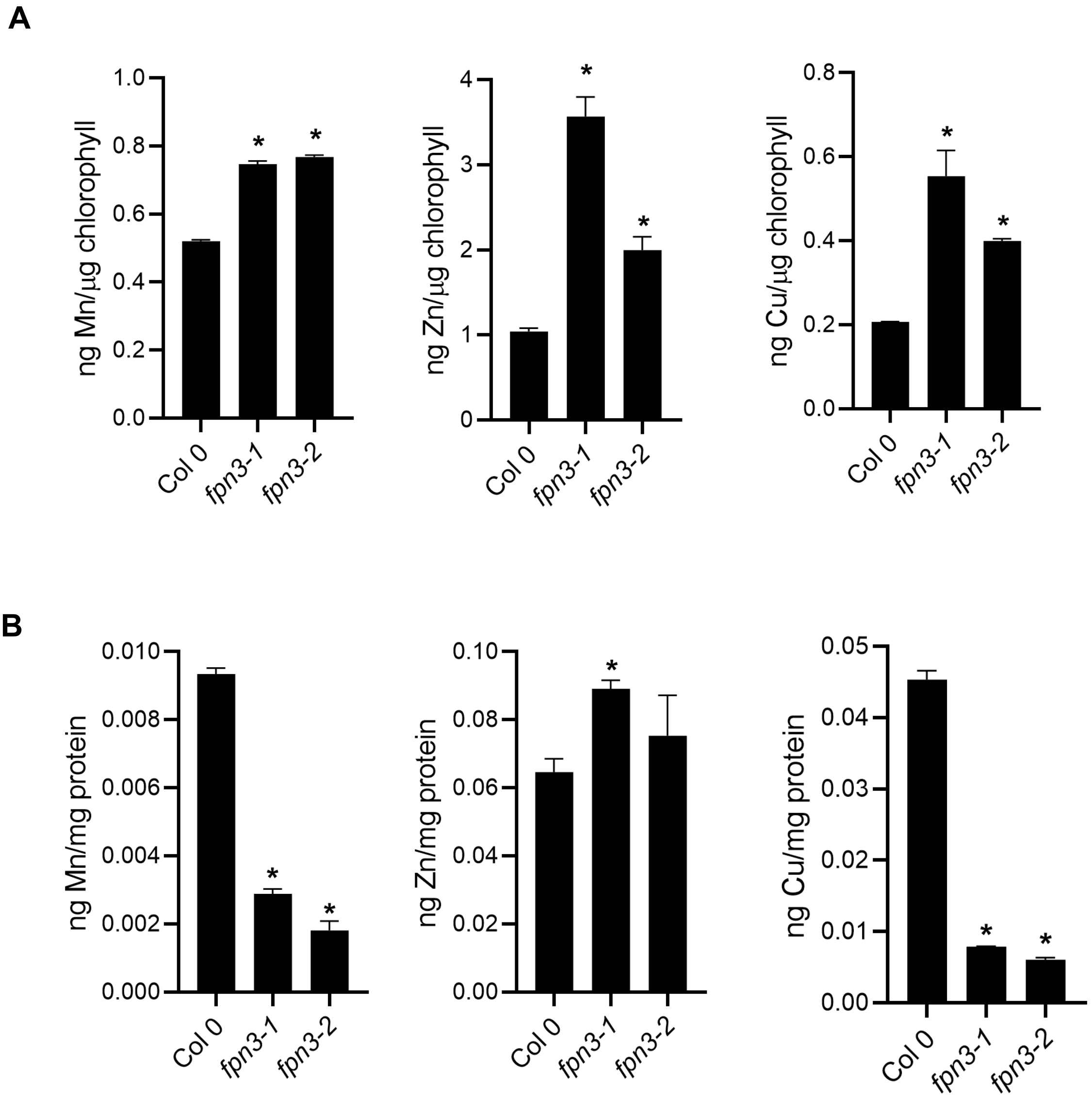


**Supplementary Figure 7. Manganese, zinc, and copper content in wild type and *fpn3* chloroplasts and mitochondria.**

ICP-MS was conducted with chloroplasts (A) and mitochondria (B) isolated from wild type (Col 0) and *fpn3*. (A) Metal content of chloroplast samples normalized with chlorophyll content. (B) Metal content of mitochondrial samples normalized with total protein. Mean values of quadruplicates for chloroplast samples and triplicates for mitochondrial samples are shown with standard error. Significant differences compared to the wild type, Col 0, are denoted (One-way ANOVA; Dunnett's test; *: $p < 0.05$).

**Supplementary Information**

**Materials and Methods**

**Western blots**

Western blots were conducted to determine the purity of isolated chloroplasts. Arabidopsis shoot protein was used as a control against the isolated chloroplasts and extracted as follows: Shoots were ground in liquid nitrogen before being suspending in extraction buffer (125 mM Tris-HCl, pH 8.8; 1% SDS; 10% Glycerol; 50 mM $Na_2S_2O_5$). After spinning down, the supernatant containing shoot protein was kept at -80ºC until use. Equal amounts of Arabidopsis proteins were separated by SDS-PAGE (10% and 15%) , transferred onto PVDF membrane, and blocked overnight at 4ºC using Intercept Blocking Buffer (LI-COR, 927-70001). Primary antibodies were diluted with blocking buffer and incubated for 1 hour at room temperature. The primary antibodies and their dilutions used were: anti-plastocyanin at 1:1000 (PhytoAB, PHY0099S), anti-Pex14p at 1:10,000 (Agrisera, AS08 372), anti-isocitrate dehydrogenase (Agrisera, AS06 203A) at 1:5000. anti-aconitase 1 at 1:500 (PhytoAB, PHY1333S), and anti-actin at 1:10,000 (PhytoAB, PHY0001), Next, blots were washed with PBS-Tween 20 before incubation in secondary antibodies for 1 hour at room temperature with the following secondary antibodies: goat anti-rabbit IRDye 800CW (Li-COR) and goat anti-mouse IRDye 800CW (Li-COR). Then, blots were washed with PBS-Tween 20 before imaging with a LI-COR Odyssey CLX (CLX-1294).
