## Supplementary Table 1 for "Ferroportin 3 is a dual-targeted mitochondrial/chloroplast iron exporter necessary for iron homeostasis in Arabidopsis"

**Supplementary Table S1: Sequences of qRT-PCR primers.**

| Gene | Forward Primer | Reverse Primer |
|---|---|---|
| *ACT2* | 5’-CCAAGCTGTTCTCTCCTTGTACGC-3’ | 5’-TCACCAGAATCCAGCACAATACC-3’ |
| *FPN3* | 5’- GTGGGTTCTTTGCCAACCATGAC-3’ | 5’- TTAGGACGGTCCAGAACTCCAG-3’ |
| *FER1* | 5’- CAACGTTGCTATGAAGGGACTAGC-3’ | 5’- ACTCTTCCTCCTCTTTGGTTCTGG-3’ |
| *FER3* | 5’- AGTGTGTTTCTGAACGAACAGGTG-3’ | 5’- AGAAGCTCCTGATCGAAATGCC-3’ |
| *FER4* | 5’- CGAGTTTCTGACAGAGCAGGTG-3’ | 5’- CGTCGCAGTTGAGCCACATATTC-3’ |
| *FRO3* | 5’- CCATACCTTTGTCACCATCACTCC-3’ | 5’- ATCCAGCCTTGCTTGCCATAAG-3’ |
| *NRAMP3* | 5’- TGGCTCTGAGCTTCTCATTGGG-3’ | 5’- TGCAACCCACAACTCCAACTGC-3’ |
