## Supplementary Table 2 for "Ferroportin 3 is a dual-targeted mitochondrial/chloroplast iron exporter necessary for iron homeostasis in Arabidopsis"

**Supplementary Table 2. NanoString Custom CodeSet**

| **Gene** | Accession | Target Sequence |
|---|---|---|
| ***ACT2*** | NM_112764.4 | TGATGAGGCAGGTCCAGGAATCGTTCACAGAAAATGTTTCTAAGCT[illegible] |
| ***BTS*** | NM_112713.4 | ATGTAATGCTTGTGCGTCTGTTCCATGCACGAGTAGGAGTACAAAAT |
| ***FER1*** | NM_120238.4 | CAAATCTCAGGCGGCTCAACACTATCCTCTTCTACATTTTCCAACGA[illegible] |
| ***FER3*** | NM_115467.4 | GTGTGGAAGAACGAGAGCACGCTGAGCTGTTGATGGAGTATCAGAA |
| ***FER4*** | NM_129588.4 | TAAAAGCTCAACCACCGACGCGCTAAGCGGCGTTGTCTTCGAGCCG[illegible] |
| ***FRO2*** | NM_001331284.1 | AAGCTTTGGTTATGGTGTGCGGAGGAAGCGGGATAACTCCGTTTAT[illegible] |
| ***FRO3*** | NM_102150.4 | CCATACCTTTGTCACCATCACTCCTCAATCACTTCCAACCGACGGAG[illegible] |
| ***FRO7*** | NM_124352.3 | CTGGATCACTCGGTGATCAAATCCTCACAAACTGGAGAGCAATCTC[illegible] |
| ***FRO8*** | NM_124395.5 | TCATGCCGGTGATCGTCACTTCTACTGGGTCCTCCCAGGCATGTTTC[illegible] |
| ***IRT1*** | NM_118089.4 | GGTCATGGTCATGGTCACGGCCCCGCAAATGATGTTACCTTACCAAT |
| ***NRAMP4*** | NM_126133.4 | ACGGTGTTTGCCAAGTCGTTTTACGGGACAGAGATAGCGGACACCA[illegible] |
| ***OPT3*** | NM_117732.7 | ATAATAGGGCAGTTCATTATCGGTTATATCCTGCCTGGAAAACCCAT |
| ***PYE*** | NM_114632.4 | ATCAGTCGAAACCTGACTTGAACACCTCTCCTGCACCCGAGTACCAT |
| ***UBC*** | NM_101307.5 | GCGTGATGTTGTTGAGCAAAGCTGGACTGCTGACTAGTAGTAGTTT[illegible] |
